# An intein-split transactivator for intersectional neural imaging and optogenetic manipulation

**DOI:** 10.1101/2021.04.05.438407

**Authors:** Hao-Shan Chen, Xiao-Long Zhang, Rong-Rong Yang, Guang-Ling Wang, Xin-Yue Zhu, Yuan-Fang Xu, Dan-Yang Wang, Na Zhang, Shou Qiu, Li-Jie Zhan, Zhi-Ming Shen, Xiao-Hong Xu, Gang Long, Chun Xu

## Abstract

The complexity of brain circuitry is manifested by numerous cell types based on genetic marker, location and neural connectivity. Cell-type specific recording and manipulation is essential to disentangle causal neural mechanisms in physiology and behavior; however, many current approaches are largely limited by number of intersectional features, incompatibility of common effectors and insufficient gene expression. To tackle these limitations, we devise an intein-based intersectional synthesis of transactivator (IBIST) to selectively control gene expression of common effectors in specific cell types defined by a combination of multiple features. We validate the specificity and sufficiency of IBIST to control common effectors including fluorophores, optogenetic opsins and Ca^2+^ indicators in various intersectional conditions *in vivo*. Using IBIST-based Ca^2+^ imaging, we show that the IBIST can intersect up to five features, and that hippocampal cells tune differently to distinct emotional valences depending on the pattern of projection targets. Collectively, the IBIST multiplexes the capability to intersect cell-type features and is compatible with common effectors to effectively control gene expression, monitor and manipulate neural activities.

## Introduction

The cell-type diversity is a hallmark of the brain circuitry that enables us to process sensory stimuli in the environment, generate appropriate behaviors and emotions and gain the knowledge from learned experiences. The properties of neural cell types play pivotal roles in governing the neural functions of circuit networks across brain regions ^1–3^, thus it becomes increasingly important for functional studies to specifically label a homogenous cell type depending on multiple features such as molecular markers and neuronal connections ^3–5^. Genetic mouse lines offer great tools to intersectionally label specific neuronal population ^6, 7^, but are often hindered by the requirement of high-level gene expression for optogenetic opsins and Ca^2+^ indicators. The breeding and crossing between multiple mouse lines can be laborious and time consuming, and region-specific control is also challenging. Combining the adeno-associated virus (AAV) vectors with transgenic mice brings powerful and convenient strategies to label a defined population in an intersectional manner ^5^ and drives sufficient expression of opsins and Ca^2+^ indicators for optogenetic manipulation and neural recording, respectively ^8^. Pioneering works have constructed AAV effectors for fluorophore and Channelrhodopsin-2 (ChR2) that selectively respond to specific combinations of particular controllers such as Cre and Flpo ^4, 9, 10^. While more intersectional effectors are made available for optogenetic opsins and Ca^2+^ indicators to match specific combinations of up to three controllers ^11^, the number of intersectional controllers/features is still limited. As the effectors for opsins and Ca^2+^ indicators require functional validations in various experimental conditions owing to the high-demand but varying expression level, it is thus desirable to fulfill both the dependency on more controllers/features and the compatibility with common effectors that could be well validated across brain regions and animal species.

To this end, we took a strategy of reconstituting a single controller to achieve the intersectional control ^7, 12^. This strategy supports multi-feature dependent reconstitution of the single controller and in turn enables the compatibility with common effectors, which could be well characterized and have reliable and sufficient gene expressions for recording or manipulation. Because most existing cell-type specific driver lines are controlled by Cre ^13, 14^, we sought to reconstitute non-Cre controllers which can be complementary for Cre, and then exploited the application with AAV and rabies vectors which are broadly used in neural circuit studies. The tetracycline-controlled transactivator (tTA) is such a widely used controller and has additional features with temporal and reversible regulation. The tTA is formed by fusing tetracycline repressor protein with an activation domain of virion protein 16 (VP16) from herpes simplex virus ^15, 16^. We reasoned that splitting tTA into these two parts would achieve separable expressions in distinct constructs, which most likely result in reconstitution of fully functional tTA. To achieve the efficient reconstitution of tTA, we took advantage of the intervening protein sequence (intein) mediated trans splicing technique, which has been shown to facilitate successful reconstitution of two parts together at the protein level in *Caenorhabditis elegans* ^17^, neural stem cell ^18^, cell line ^19^ and mouse *in vivo* ^7^. Compared to other reconstitution approaches such as leucine zipper mediated dimerization and adapter pair based covalent bonding, the intein approach introduces minimal heterologous sequence and exhibits better kinetics and higher activity recovery ^17, 20–22^.

In summary, we developed a system with intein-based intersectional synthesis of transactivator (IBIST) to achieve the intersectional expression of a single controller. It in turn fulfills the compatibility with common effectors and multiplexes the intersectional features based on other existing controllers. This system supports: (1) flexible design of modular expression of split tTA in various conditions, (2) compatibility with common effectors dependent on tetracycline-responsive promoter element (TRE), and (3) versatile use with the viral vectors and rich resources of increasing mouse driver lines ^5, 13, 14, 23^. To demonstrate the capability of IBIST for intersectionally expressing opsin and Ca^2+^ indicators, we created various AAV and rabies vectors to express tTA fragments controlled by site-specific recombinase, promoters or circuit tracing. We validated the specificity and efficiency of IBIST tools both *in vitro* and *in vivo*, and then performed electrophysiological and photometric recording to demonstrate the successful application of IBIST tools for optogenetics and Ca^2+^ imaging.

## Results

### The design of IBIST and the validation *in vitro* and *in vivo*

We split the tTA into N-terminal (Tet repressor, TetR) and C-terminal (three tandem minimal VP16 activation domains) parts from its structure junction ^16^. The N-terminal part was further tagged with a blue fluorescence protein (BFP, Fig.1A). To facilitate the tTA reconstitution from two split parts, we attached the N- and C-fragment of intein sequences of Gp41-1 ^17, 20^ to the N- and C-terminal parts of tTA because these intein sequences were reported to be more efficient and faster for protein splicing compared to other intein sequences ^24, 25^. This pair of ready-to-reconstitute subunits are referred as tTAN and tTAC respectively and are designed to form functional tTA after intein-mediate splicing (Fig.1 A-C).

**Figure 1.**
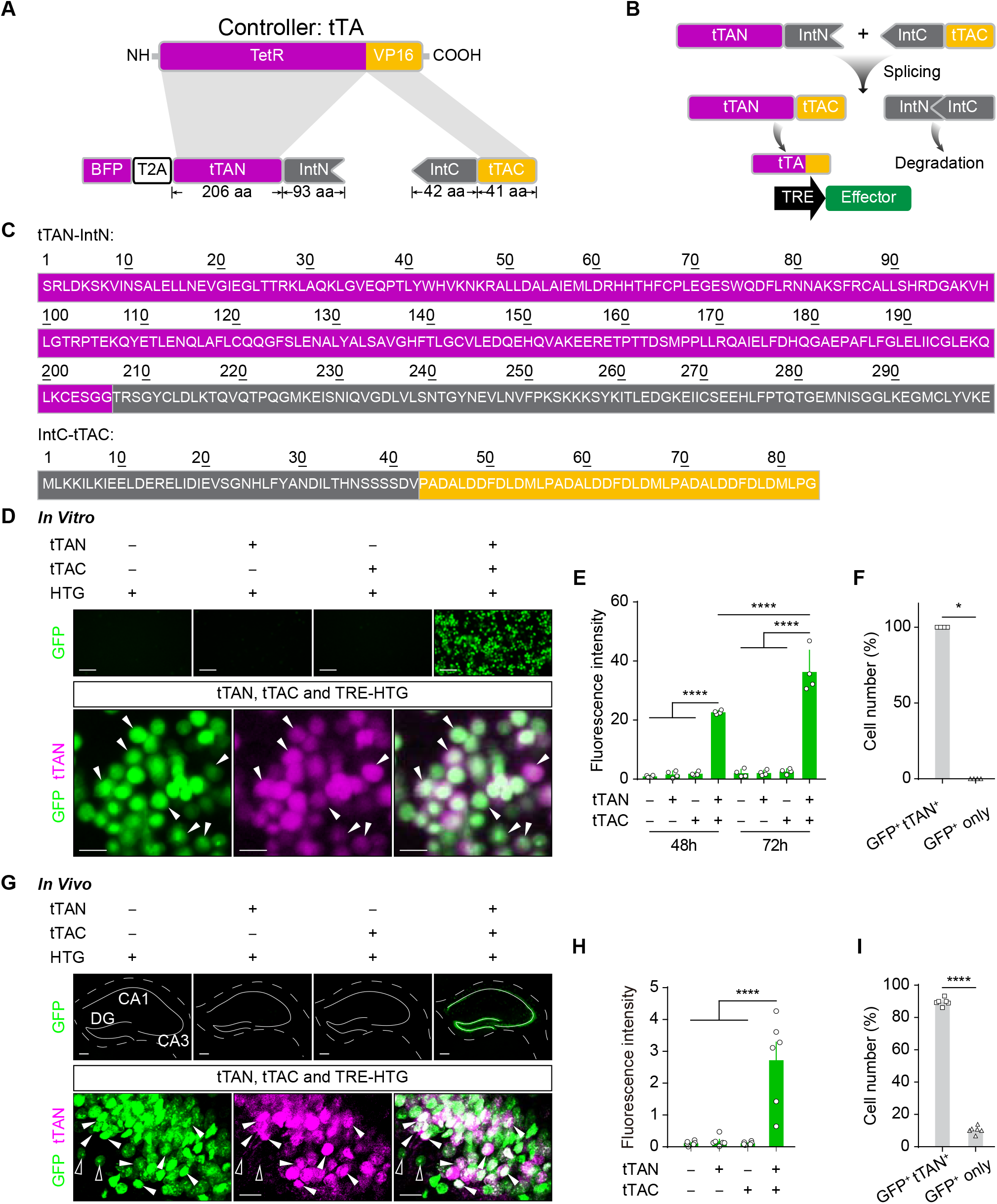
The design of IBIST and the *in vitro* and *in vivo* validation. **A**, Scheme depicting construct design of intein-split tTA. **B**, Diagram depicting the reconstitution of tTA by intein-based protein splicing. **C**, Amino acid sequence of intein-fused tTA fragments. **D**, *Top*, examples of green fluorescence in HEK293T cells 72 h after transfections of reporter plasmid (TRE-HTG) and intein-split tTA plasmids with EF1α promoter (tTAN, tTAC, both or neither). Scale bars: 20 μm. *Bottom*, examples showing the fluorescence of HTG reporter (shown in green) and tTAN (BFP tagged, shown in magenta) in HEK293T cells with all three plasmids transfected. Arrows: examples of co-labeled cells. Scale bars: 5 μm. **E**, Summary of green fluorescence intensity of HEK293T cells (**D**, top) at 48 h and 72 h post transfection (n = 4 FOV in each group). One-way ANOVA revealed a significant difference in green fluorescence intensities between groups (F_(7, 24)_ = 95.73, P < 0.0001). Tukey’s multiple comparisons test revealed that fluorescence intensity in tTAC+tTAN group at 72 h is significantly higher than that at 48 h post transfection (P <0.0001), and both are significantly higher than other groups at the same time post transfection (P < 0.0001 for all). **F**, Percentage summary of cells labeled by HTG and tTAN in **D** (100±0 co-labeled vs. 0±0 reporter only; Mann-Whitney U test, P < 0.05, n = 4 FOV). **G**, *Top*, examples showing reporter fluorescence (AAV-TRE-HTG) after its injection into hippocampus and co-injections of AAVs (CaMKIIα promoter) for tTAN, tTAC, both or neither, respectively. Scale bars: 200 μm. *Bottom*, examples showing fluorescence of tTAN (shown in magenta) and HTG reporter (shown in green) in hippocampal cells. Open arrows: reporter only. Filled arrows: co-labeled. Scale bars: 20 μm. **H**, Summary of green fluorescence intensities in hippocampus (Each group, n = 6 FOV, N = 3 animals). One-way ANOVA revealed significant fluorescence differences between groups (F_(3, 20)_ = 20.57, P < 0.0001) and Turkey’s multiple comparisons revealed that the fluorescence is significantly higher in TRE-HTG+tTAC+tTAN group than in others (P < 0.0001 for all). **I**, Percentage summary of cells labeled by reporter and tTAN (89.6±0.9% co-labeled vs. 10.4±0.9% reporter only; paired *t*-test, P < 0.0001, n = 6 FOV, N = 3 animals).

To validate whether the tTA could be successfully reconstructed in mammalian cells, we expressed tTAN and tTAC as well as a TRE controlled GFP reporter ^26^ in cultured HEK293T cells (see methods; TRE-dependent expression of histone-GFP, TVA and rabies glycoprotein, termed TRE-HTG). The fluorescence and protein level of TRE-dependent reporter became prominent after transfection, which was in stark contrast to that in control groups (Fig. 1D, E and Fig. S1A). These results confirmed that split tTA are specifically rescued in the targeted cells with the help of intein-mediated splicing. All GFP positive cells also expressed the fluorescence tag of tTAN (100±0%, n = 4 field of view, termed FOV; Fig. 1D and F), suggesting that the tTA reconstitution is highly specific.

To test whether similar strategy is applicable for reverse tTA (rtTA), we engineered the constructs for split rtTA with same intein sequences (Fig. S1B). We first transfected the HEK293T cells with plasmids for C-terminal part of rtTA (rtTAC) and reporter TRE-mCherry. To test whether the rtTA fragments can be expressed by viral vectors for neural tracing, we then inoculated the cells with a rabies variant expressing the N-terminal part of rtTA and the GFP tag (termed RV-rtTAN), which could be a retrograde transsyanptic tracer ^27^. In the presence of doxycycline, the fluorescence and protein level of TRE-mCherry became prominent after transfection, but nondetectable in the control group (Fig. S1A, C and E). Nearly all of mCherry positive cells were co-labeled by the RV-rtTAN (96.4±1.5% co-labeled vs. 3.6±1.5% reporter only; paired *t*-test, P < 0.0001, n = 4 FOV; Fig. S1D and F), suggesting the rtTA reconstitution is highly specific. Taken together, the fragments of tTA and rtTA are both able to efficiently and specifically reconstitute tTA and rtTA in the mammalian cells with the help of intein sequences, respectively.

To validate whether the split tTA could form functional reconstitution *in vivo*, we produced the AAV vectors and injected into hippocampus with the AAVs of TRE-HTG and CaMKIIα dependent tTAC and BFP tagged tTAN. We observed strong fluorescence expression of reporter AAV-TRE-HTG in the hippocampus, whereas little fluorescence was seen in the control animals (Fig. 1G and H). Majority of HTG-positive cells co-expressed endogenous fluorescence tag of tTAN (89.6±0.9% co-labeled vs. 10.4±0.9% reporter only; paired *t*-test, P < 0.0001, n = 6 FOV; Fig. 1G and I). Together, these results suggest that intein-split tTA/rtTA fragments are able to be specifically reconstructed in an intersectional manner.

### The function of IBIST-controlled optogenetic manipulations and Ca^2+^ imaging

Combining intersectional approaches with optogenetic manipulations is instrumental to delineate neural circuit functions ^8^. Although many intersectional labeling strategies exist for fluorophore labeling, it remains to be tested whether these strategies are applicable for optogenetic opsins because high expression of opsin is required for optogenetic manipulations in functional studies. We sought to test this for the IBIST tools by injecting hippocampus with AAVs expressing TRE-oChiEF-mCherry ^28^ and CaMKIIα-dependent tTAN and tTAC, respectively (Fig. 2A). As expected, potent mCherry fluorescence were only seen in animals with all three AAVs injected compared to control animals with AAV-tTAN or AAV-tTAC omitted (Fig. 2B). To further validate the faithfulness of reporter expression, we carried out immunostaining to amplify the endogenous fluorescence. We found that nearly all TRE-oChiEF positive cells were also co-labeled by tTAN tag (96.2±1.6% co-labeled vs. 3.8±1.6% reporter only; paired *t*-test, P < 0.0001, n = 6 FOV; Fig. S2A and B).

**Figure 2.**
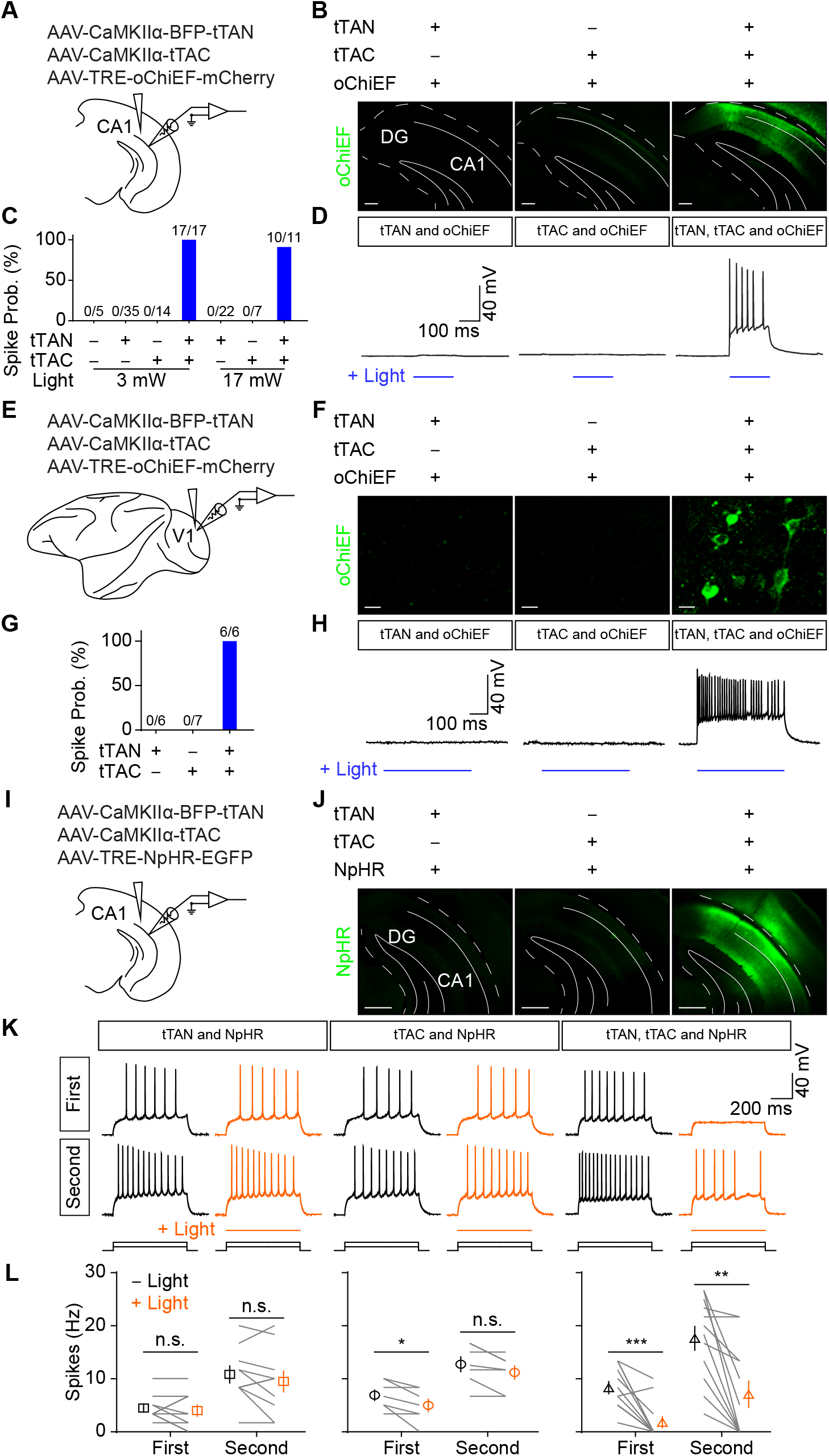
The IBIST-based optogenetic manipulation. **A**, Scheme depicting AAV injections into hippocampus. **B**, Examples showing mCherry fluorescence (shown in green) in hippocampus injected with AAV-TRE-oChiEF and AAVs (CaMKIIα promoter) of tTAN, tTAC or both, respectively. Scale bars: 200 μm. **C,** Summary of spike probabilities (prob.) in hippocampal cells evoked by whole-field blue LED light (Left: 3 mW at the objective; Right: 17mW at the objective) from animals injected with AAV-TRE-oChiEF and other AAVs (CaMKIIα promoter): none, N = 1 animal; tTAN, N = 6 animals; tTAC, N = 2 animals; tTAN+tTAC, N = 4 animals. **D**, Examples showing whole-cell current-clamp recording from hippocampal cells in brain slices with blue LED light stimulation (blue line). All animals were injected with AAV-TRE-oChiEF and co-injected with AAVs (CaMKIIα promoter) of either tTAN, tTAC or both. **E**, Scheme depicting AAV injections into multiple regions in V1 of *Macaca fascicularis*. **F**, Examples showing mCherry fluorescence (shown in green) in V1 brain region after injected with AAV-TRE-oChiEF-mCherry and AAVs (CaMKIIα promoter) of tTAN, tTAC or both, respectively. Scale bars: 10 μm. **G**, Summary of spike probabilities (prob.) evoked by whole-field blue LED light (3 mW) from animals injected with AAV-TRE-oChiEF and other AAVs. **H**, Examples showing whole-cell current-clamp recording in brain slices with blue LED light stimulation (blue line). All groups were injected with AAV-TRE-oChiEF and co-injected with AAVs (CaMKIIα promoter) of either tTAN, tTAC or both. **I**, Scheme depicting AAV injections into hippocampus. **J**, Examples showing GFP fluorescence in hippocampus after injected with AAV-TRE-NpHR and AAVs (CaMKIIα promoter) of tTAN, tTAC or both, respectively. Scale bars: 500 μm. **K**, Examples showing whole-cell current-clamp recording from hippocampal cells in brain slices upon current injections with 40 pA steps (first and second sweeps with prominent spikes; yellow with light; black without light). **L,** Frequency of spikes in hippocampal cells in the absence (black) or presence (yellow) of 589 nm laser light (18 mW) from animals injected with AAV-TRE-NpHR and other AAVs (CaMKIIα promoter, N = 2 animals for each group). The tTAN+tTAC animals showed a significant decrease in frequency of spikes after light stimulation (first sweep, 8.3±1.1 Hz vs. 1.8±1.0 Hz with light; paired *t*-test, P = 0.0004; second sweep, 17.6±2.3 Hz vs. 7.1±2.5 Hz, paired *t*-test, P = 0.0012). The tTAN animals: first sweep, 4.5± 0.7 Hz vs. 4.0± 1.1 Hz with light; paired *t*-test, P = 0.5414; second sweep, 10.8±1.6 Hz vs. 9.5±2.0 Hz with light; paired *t*-test, P = 0.21. The tTAC animals: first sweep, 6.9±1.0 Hz vs. 5.0±1.2 Hz with light; paired *t*-test, P = 0.03; second sweep, 12.7±1.4 Hz vs. 11.2±1.2 Hz with light; paired *t*-test, P = 0.0523).

Using patch-clamp recording with optogenetic stimuli in acute brain slice (see methods), we found that the blue LED light robustly evoked spikes in hippocampal neurons from the AAV infected brain slices (200 ms constant light stimuli: 100%, 17/17 cells, Fig. 2C and D; 2 ms light pulses at 25 Hz; Fig. S2C). In control animals injected with AAVs of tTAN or tTAC omitted, no spikes were evoked by the blue LED light (tTAN, 0/35 cells; tTAC, 0/14 cells; Fig. 2C and D). When we set the light stimuli at the maximal power of the LED device (17 mW at the 60X objective), we still observed no spikes in hippocampal cells from control animals (tTAN, 0/22 cells; tTAC, 0/7 cells; Fig. 2C).

The viral vectors serve as indispensable tools in primate research as transgenic primate animals are very limited. Thus, we further tested whether the same viral vectors were able to reconstitute tTA in primate brain. We injected same AAVs into primate visual cortex and performed patch-clamp recording with optogenetic stimuli (Fig. 2E). The fluorescence tag of effector (shown in green) was only seen in brain slices with all three AAV vectors injected but not in brain slices with control injections (Fig. 2F). Accordingly, low-intensity LED light (3 mW) successfully elicited spikes in all fluorescent cells but not in cells from slices with control injections (tTAN, 0%, 0/6 cells; tTAC, 0%, 0/7 cells; tTAC+tTAN, 100%, 6/6 cells; Fig. 2G and H).

Next, we validated the optogenetic inhibition by Halorhodopsin (NpHR) ^29^ controlled by reconstituted tTA (Fig. 2I and J). In the acute brain slices from mice injected with all three AAV vectors, the yellow light was able to effectively inhibit neuronal spikes evoked by current injections (first sweep, 8.3±1.1 Hz vs. 1.8±1.0 Hz with light; paired *t*-test, P = 0.0004; second sweep, 17.6±2.3 Hz vs. 7.1±2.5 Hz, paired *t*-test, P = 0.0012; Fig. 2K and L). In control slices from animals with either AAV-tTAN or AAV-tTAC omitted, the yellow light did not exert obvious changes in the neuronal spiking (The tTAN animals: first sweep, 4.5±0.7 Hz vs. 4.0±1.1 Hz with light; paired *t*-test, P = 0.5414; second sweep, 10.8 ± 1.6 Hz vs. 9.5±2.0 Hz with light; paired *t*-test, P = 0.21. The tTAC animals: first sweep, 6.9±1.0 Hz vs. 5.0 ± 1.2 Hz with light; paired *t*-test, P = 0.03; second sweep, 12.7±1.4 Hz vs. 11.2±1.2 Hz with light; paired *t*-test, P = 0.0523; Fig. 2K and L). These results suggest that the reconstituted tTA is capable of driving sufficient and specific expression of optogenetic opsin for neuronal manipulations in both mice and primates.

Monitoring specific population of neurons defined by intersectional conditions is extremely useful to dissect neuronal functions but also requires high expression of Ca^2+^ indicators *in vivo*. To test whether the IBIST is capable of driving sufficient and specific expression of Ca^2+^ indicators such as GCaMP6s ^30^, we injected into hippocampus with AAVs of CaMKIIα-tTAN, CaMKIIα-tTAC and TRE-GCaMP6s. After implanting with optical fiber above the hippocampus, we performed photometric recording of Ca^2+^ activity in the hippocampus while the animal was exploring in the fear conditioning context (Fig. S3A). When the animal received several foot shocks, we observed that strong Ca^2+^ activity was evoked by the foot shocks whereas no detectable Ca^2+^ signal was recorded in control animals with either AAV-CaMKIIα-tTAN or AAV-CaMKIIα-tTAC omitted (Fig. S3B-D). Interestingly, we found that the shock-evoked Ca^2+^ activity was further enhanced in the second conditioning session, indicating that these hippocampal neurons became sensitized to the shock stimuli. Importantly, the Ca^2+^ signal was still not detectable in the control animals (Fig. S3D). Taken together, these results confirmed that IBIST is able to drive specific and sufficient effector expression for optogenetic opsins and Ca^2+^ indicators *in vivo*.

### IBIST-based tools to define cells by genetic marker and neural connectivity

The genetic marker, neural connectivity and neuronal location constitute the major features to define the cell type. While neuronal locations are conveniently controlled by stereotaxic injections of viral vectors, we sought to test the compatibility of IBISIT-based tools when defining the other two types of features in various means. We first tested whether IBIST tools were able to combine features of anterograde labeling and promoter-based viral labeling. The hippocampal dorsal CA3 (dCA3) cells send collaterals to both pyramidal cells and interneurons in ventral CA1 (vCA1). We designed a strategy to combine AAV serotype 1 mediated anterograde transsyanptic tracing ^31^ with pyramidal-cell-specific fluorophore labeling in vCA1, aiming to specifically label pyramidal cells with direct inputs from dCA3. We injected AAV1-EF1α-tTAN into the dCA3 and injected into vCA1 with AAV-CaMKIIα-tTAC and reporter AAV-TRE-HTG (Fig. S4A). Although AAV1-EF1α-tTAN travelled in both retrograde and anterograde directions and in turn labeled cells in DG, CA1 and subiculum, we observed specific labelling of CA3-targeted CA1 pyramidal cells owing to the local infection of AAV-CaMKIIα-tTAC in vCA1 (Fig. S4B). As comparison, no GFP expression was detected in CA1 from the control animals. We then tested whether IBIST tools were able to combine features of anterograde labeling and Cre-dependent labeling in transgenic mice. We used AAV1-Flpo as an anterograde tracer and combined injections of AAV-FRT-tTAN-BFP, AAV-DIO-tTAC, AAV-DIO-mCherry and AAV-TRE-GCamP6s in *SOM-ires-Cre* mice ^*13*^, aiming to specifically label SOM+ cells with direct inputs from dCA3 (Fig. S4C). As a result, we observed largely specific labeling of CA3-targeted SOM+ cells (66.9±2.7% co-labeled by GCa. and mCherry vs. 33.1±2.7% GCa. only; paired *t*-test, P = 0.0004, n = 8 FOV; Fig. S4D and E). We further tested whether IBIST tools are able to combine features of retrograde labeling and promoter-based viral labeling. Using tTAN-expressing rabies vectors, we selectively labeled ventral hippocampus upstreams (CA3 cells) with TRE reporter (96.3±0.4% co-labeled vs. 3.7±0.4% reporter only; paired *t*-test, P < 0.0001, n = 4 FOV; Fig. S4F-H). We continued to test whether the tTA reconstitution is feasible with fragments controlled by recombinase in transgenic animals, we engineered Flpo-dependent construct for tTAN and Cre-dependent construct for tTAC, and injected AAVs in hypothalamus where cell types are typically defined by multiple molecular markers (Fig. S4I). In the transgenic mice of *Vgat-Flpo*/*V*g*lut2-Cre*, we indeed observed specific reporter expression when the tTAN and tTAC were expressed in Cre and Flpo dependent manners, respectively (Fig. S4J). In situ hybridization revealed that most reporter-expressing cells were positive for conditional marker *Vgat* (78.0±4.6%; n = 10 FOV; Fig. S4K and L). These results demonstrated that IBIST-based tools are capable of defining cell types with dual features by various means using viral vectors and transgenic animals.

To validate whether these dual-feature cell types have sufficient gene expression for effectors such as Ca^2+^ indicators and optogenetic opsins, we defined vCA1 cells by the genetic marker and their connections with dCA3. To specifically image vCA1 excitatory neurons with direct inputs from dCA3, we injected AAV1-EF1α-Flpo into dCA3 and injected into vCA1 with AAV-FRT-BFP-tTAN, AAV-CaMKIIα-tTAC and effector AAV-TRE-GCaMP6s (Fig. 3A and B). We performed photometric recording from those excitatory neurons in vCA1 while animals underwent open field test and contextual fear conditioning (Fig. 3C). We observed significant increase in Ca^2+^ signals when animals were entering the center of the open field, which were not seen in control animals (tTAN, 0.4±0.2, N = 6 animals; tTAC, 0.5±0.2, N = 7 animals; tTAN+tTAC, 2.0±0.6, N = 4 animals; F_(2,14)_ = 8.113, P = 0.0046; Turkey’s multiple comparisons, P < 0.01; Fig. 3D and E). These results indicated that these CA3-targeted CA1 cells exhibit anxiety-related signals. When animals received foot shocks, significant Ca^2+^ signals were detected as well (First conditioning: tTAN, 0.3±0.2%, N = 6 animals; tTAC, 0.3±0.1%, N = 7 animals; tTAN+tTAC, 4.1±0.8%, N = 4 animals; F_(2,14)_ = 31.79, P < 0.0001; Turkey’s multiple comparisons, P < 0.0001. Second conditioning: tTAN, 0.4±0.1%, N = 6 animals; tTAC, 1.0±0.3%, N = 7 animals; tTAN+tTAC, 5.7±0.5%, N = 4 animals; F_(2,14)_ = 67.15, P < 0.0001; Turkey’s multiple comparisons, P < 0.0001; Fig. 3F and G). Next, we adopted similar IBIST-based strategy to enable specific ChR2 expression in pyramidal cells with dCA3 inputs (Fig. S5A-D). The blue light successfully evoked spikes in vCA1 neurons with direct inputs from dCA3 (33%, 3/9 cells, 3 mW; 56%, 5/9 cells, 17 mW; Fig. S5E and F). Finally, we utilized different sets of viral vectors in *SOM-ires-cre* mice to enable specific NpHR expression in SOM+ cells with dCA3 inputs (Fig. 5G and H). As expected, the yellow light effectively cracked down the neuronal spikes (First sweep, 9.6±1.4 Hz vs. 2.0±1.0 Hz with light; Wilcoxon matched-paired signed rank test, P = 0.0313; Second sweep, 28.2±4.3 Hz vs. 14.7±4.2 Hz with light; Wilcoxon matched-paired signed rank test, P = 0.0313; Fig. S5I and J). These results confirmed that the IBIST tools enabled dual-feature cells with specific and sufficient expression of Ca^2+^ indicators and optogenetic opsins.

**Figure 3.**
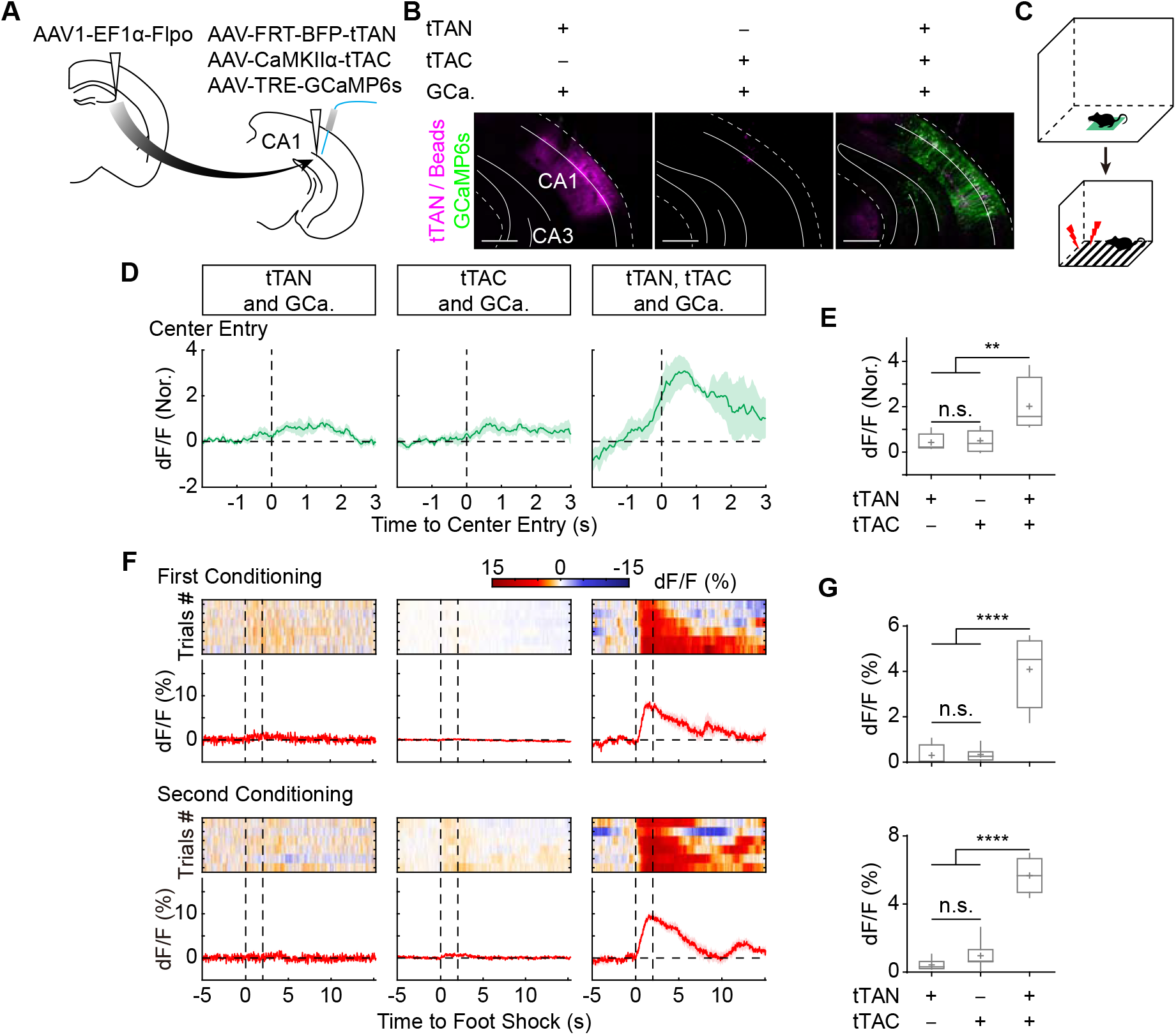
The IBIST-based Ca^2+^ recording for cells defined by genetic marker and neural connectivity. **A**, Scheme illustrating AAV injections to define cells by two features: anterograde transsynaptic tracing and pyramidal-cell-specific (CaMKIIα promoter) fluorophore labeling. **B**, Examples showing fluorescence of the ventral hippocampus from animal injected with various combinations of AAVs of Flpo, tTAN (shown in magenta), tTAC and TRE-GCaMP6s (GCa., shown in green) and co-injected with blue beads (shown in magenta). Scale bars: 500 μm. **C**, Scheme illustrating the behavioral experiments. **D**, Examples of averaged traces of Ca^2+^ signals upon center entry in the open field test. **E**, Box plot of normalized Ca^2+^ responses upon center entry. One-way ANOVA revealed significant differences between groups (tTAN, 0.4±0.2, N = 6 animals; tTAC, 0.5±0.2, N = 7 animals; tTAN+tTAC, 2.0±0.6, N = 4 animals; F_(2,14)_ = 8.113, P = 0.0046; Turkey’s multiple comparisons, P < 0.01). **F,** Examples of heatmaps and averaged traces of Ca^2+^ signals upon foot shocks (between vertical dash lines, 6 trials) in the first and second fear conditioning session. **G,** Box plot of Ca^2+^ responses. One-way ANOVA revealed significant differences between groups in first conditioning (tTAN, 0.3±0.2%, N = 6 animals; tTAC, 0.3±0.1%, N = 7 animals; tTAN+tTAC, 4.1±0.8%, N = 4 animals; F_(2,14)_ = 31.79, P < 0.0001; Turkey’s multiple comparisons, P < 0.0001) and second conditioning (tTAN, 0.4±0.1%, N = 6 animals; tTAC, 1.0±0.3%, N = 7 animals; tTAN+tTAC, 5.7± 0.5%, N = 4 animals; F_(2,14)_ = 67.15, P < 0.0001; Turkey’s multiple comparisons, P < 0.0001).

In summary, both tTAN and tTAC are compatible with viral vectors for anterograde and retrograde circuit tracing and controllable by recombinase such as Cre and Flpo. Therefore, these IBIST tools are able to define cell types with two types of features by various means, achieve cell-type specific fluorophore expression and suffice for Ca^2+^ imaging and optogenetic manipulation.

### IBIST-based Ca^2+^ imaging for cells with multiple features

To exploit the application of IBIST tools to label cells with multiple projections, we turned to ventral hippocampus where neuronal functions diversify at the single-cell level by their distinct patterns of downstream connections and functional roles in fear conditioning, social memory and drug-induced place preference ^32–36^. While medial prefrontal cortex (mPFC) and amygdala projection cells exhibited distinct functions compared to single-projection cells ^37^, it is important and interesting to investigate how different subgroups of vCA1 projection cells respond to external stimuli with distinct emotional valences. To address this question, we used IBIST tools to label vCA1 cells with triple features by two projection targets and pyramidal cell specific CaMKIIα promoter. We injected retrograde AAVs (AAVretro) of CaMKIIα-tTAN and CaMKIIα-tTAC into one or two vCA1 downstream targets including amygdala, nucleus accumbens (NAc) and mPFC respectively, and injected AAV-TRE-GCaMP6s into vCA1 (Fig. 4A). We found that the double-projection cells were successfully labeled, which were not seen in control experiments with either AAVretro-CaMKIIα-tTAN or AAVretro-CaMKIIα-tTAC omitted (Fig. 4C). Using photometric recording from the vCA1 when animals underwent conditioned place preference (CPP) training and fear conditioning (Fig. 4B, Fig. S6A-D), we observed that significant Ca^2+^ activity was evoked in most types of vCA1 projection cells by the water reward, sucrose reward and foot shocks (Fig. 4D-G). No detectable Ca^2+^ signals were recorded in the control animals (Fig. 4D-G). As shown in Fig. 4h, foot shock activated vCA1 cells in a target-dependent manner (Amy, 5.0±1.3%; NAc, 5.4±2.8%; mPFC, 0.6±0.2%; Amy and NAc, 4.1±0.9%; NAc and mPFC 3.4±0.4%; mPFC and Amy, 2.1±0.5%). The Ca^2+^ activity was highest in NAc-projecting cells, second highest in Amy-projecting cells and lowest in mPFC-projecting cells. The double-projection vCA1 cells showed moderate activities in between. The NAc-projecting and NAc-Amy-projecting vCA1 cells responded stronger to sucrose reward than other types of projection cells (Amy, 0.2±0.2%; NAc, 0.5±0.1%; mPFC, 0.1±0.04%; Amy and NAc, 0.4±0.2%; NAc and mPFC 0.2±0.1%; mPFC and Amy, 0.1±0.04%). These results suggest that the Ca^2+^ signals in response to sucrose reward in double-projection cells is dominated by one downstream target. Notably, water reward elicited very much specific activation of NAc-projecting vCA1 cells compared to other types of projecting cells (Amy, 0.02±0.1%; NAc, 0.2±0.1%; mPFC, 0.02±0.02%; Amy and NAc, 0.004±0.1%; NAc and mPFC 0.02±0.1%; mPFC and Amy, 0.03±0.1%). Interestingly, the omission of water reward elicited small inhibitory signals in many types of vCA1 projection cells (Amy, −0.3±0.1%; NAc, −0.2±0.1%; mPFC, −0.2±0.1%; Amy and NAc, −0.1±0.1%; NAc and mPFC −0.1±0.1%; mPFC and Amy, −0.02±0.1%).

**Figure 4.**
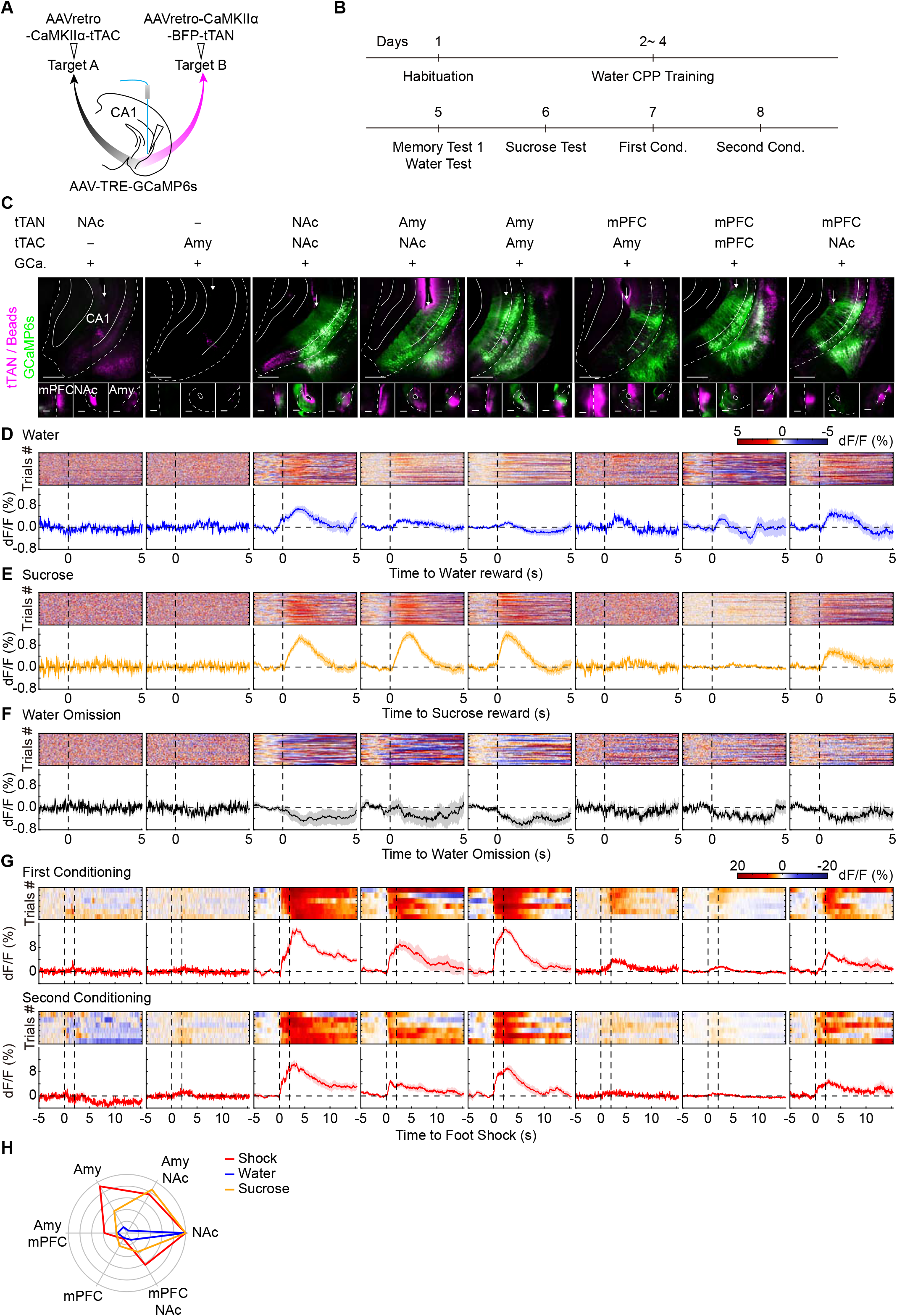
The IBIST-based Ca^2+^ recording for cells defined by multiple projections. **A**, Scheme illustrating AAV injections to define cells by three features: dual retrograde tracings and pyramidal-cell-specific (CaMKIIα promoter) Ca recording. **B**, Schematic showing the behavioral design of water CPP and fear conditioning. **C**, Examples of fluorescence images in ventral hippocampus (top, injected with AAV-TRE-GCaMP6s, shown in green) and downstream regions (bottom). AAVretro-tTAN (shown in magenta) and AAVretro-tTAC were injected into two downstream regions, co-injected into one region, or omitted. All the AAVs were co-injected with blue beads (shown in magenta). The arrows indicate optical fiber tracks. mPFC: medial prefrontal cortex. NAc: nucleus accumbens. Amy: amygdala. Scale bars: 500 μm. **D-G**, Examples of heatmaps and averaged traces of Ca^2+^ signals upon water reward (**D**, blue), sucrose reward (**E**, yellow), water omission (**F**, black) and foot shock (**G**, red). Similar photometric recording results were obtained from 3 replicates in each group except 2 replicates in the group of tTAC+tTAN in NAc. **H**, Polar summary plot of normalized Ca^2+^ signals for various types hippocampal projection cells in **C**.

Finally, we tested whether the IBITS tools could label specific types of hippocampal cells based on more downstream targets. We injected AAV-TRE-GCaMP6s into vCA1 and AAVs (AAVretro) of EF1α-FRT-tTAN, EF1α-Flpo, EF1α-DIO-tTAC, CaMKIIα-Cre into NAc, mPFC, Lateral septum (LS), and Amy, respectively. As expected, we observed prominent fluorescence of GCaMP6s in vCA1, which was absent in controls where either tTAN or tTAC was omitted from injections (Fig. 5A and B). Accordingly, we recorded robust Ca^2+^ signals from vCA1 upon foot shock (tTAN+tTAC vs. Ctrl., First conditioning, 1.6±0.5% vs. 0.2±0.04%; Mann-Whitney U test, P = 0.0238; Fig. 5C and D). Interestingly, these Ca^2+^ signals exhibited adaption over trials (First Conditioning, first 3 trials vs. last 3 trials, 2.1±0.7% vs. 1.2±0.4%; paired *t*-test, P = 0.0328). Therefore, the IBIST-based Ca^2+^ imaging is applicable for cell types defined by five features including one promoter and four different projections. Taken together, the selective recording of Ca^2+^ activity in various types of projection cells in vCA1 revealed distinct activity profiles in response to aversive and appetitive stimuli, highlighting the importance to define cell types based on multiple projection targets.

**Figure 5.**
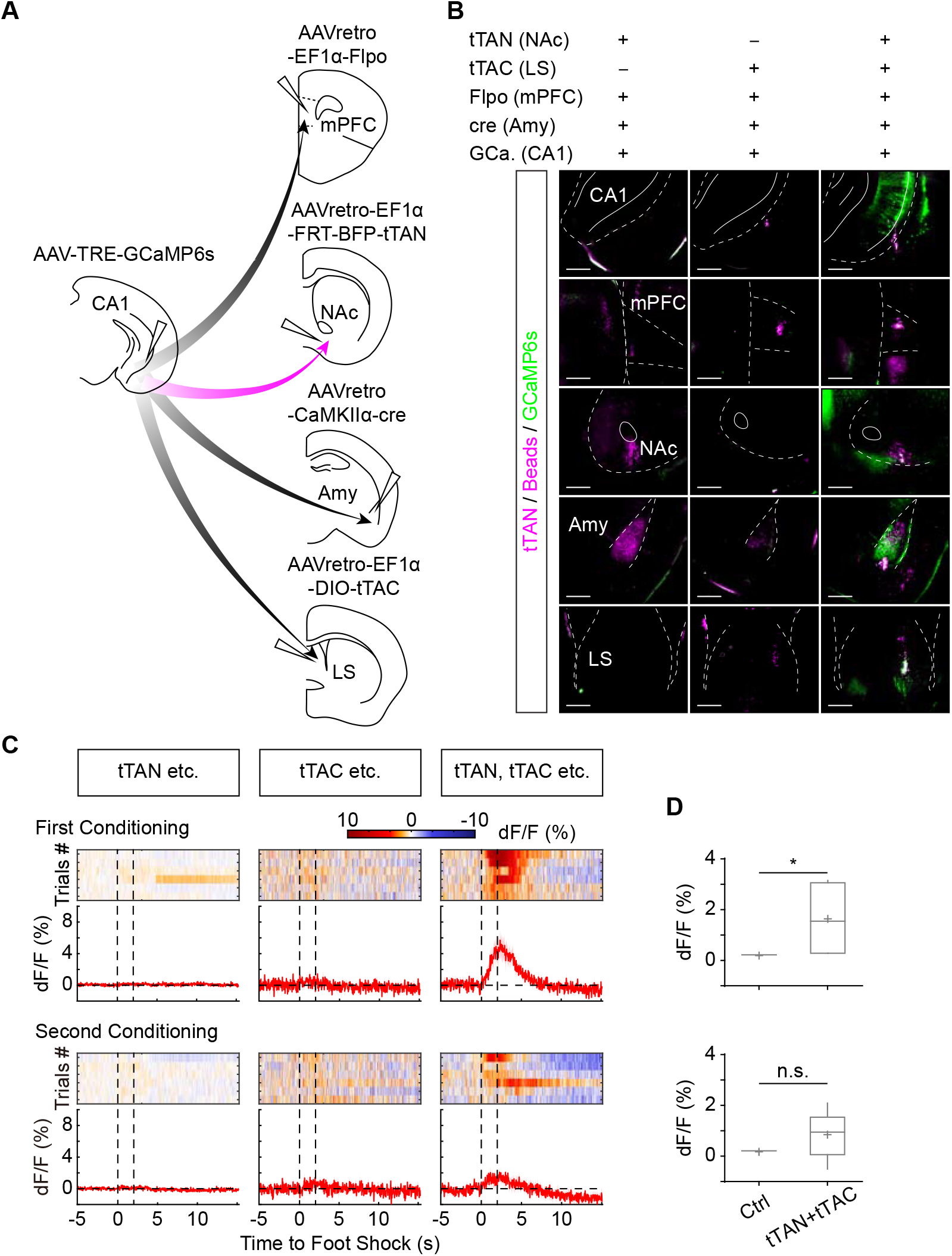
Quintuple features-based Ca^2+^ recording. **A**, Scheme illustrating AAV injections to define cells by five features including quadruple retrograde tracings and pyramidal-cell-specific (CaMKIIα promoter) Ca^2+^ recording. **B**, Examples showing fluorescence of GCaMP6s (green), tTAN (magenta) and blue beads (magenta). mPFC: medial prefrontal cortex. NAc: nucleus accumbens. Amy: amygdala. LS: lateral septum. Scale bars: 500 μ Examples of heatmaps and averaged traces of Ca^2+^ signals upon foot shocks between dash vertical lines. **D**, Box plots for foot shocks evoked Ca^2+^ responses in tTAN+tTAC (N = 6 animals) and control (1 animal for tTAN and 2 animals for tTAC) groups. First conditioning, 1.6±0.5% vs. 0.2±0.04%; Mann-Whitney U test, P = 0.0238; Second conditioning, 0.8±0.4% vs. 0.2±0.1%, Mann-Whitney U test, P = 0.2381.

## Discussion

Our work demonstrated that the IBIST can be efficiently reconstituted in various ways depending on promoter, brain regions, neuronal connections and site-specific recombinase. On one hand, the IBIST toolset is able to define the cell types by intersecting up to five features, which does more than prior tools do. In principle, the potential number of intersectional features is multiplexed by the number of fragmented tTA (Fig. S7). On the other hand, many existing TRE-based viral vectors become immediately ready-to-use for intersectional approaches, and new common effectors could be adaptable for intersectional control in a wide range of contexts.

While intersectional labeling neurons by fluorophores is widely used, it is still challenging to achieve the specificity and sufficiency for optogenetic manipulations and Ca^2+^ imaging *in vivo*. While pioneering works developed intersectional effectors for opsins and Ca^2+^ indicators ^11^, the specific combinations of controllers would require sophisticated construct design, which may suffer from insufficient gene expression in some conditions. The IBIST tools are able to resolve these concerns by intersectional synthesis of controller (tTA) which does not need that high expression level as the effector does. Accordingly, it is more straightforward to design constructs for IBIST effectors and more feasible to achieve sufficient gene expression for opsins and Ca^2+^ indicators. In our study, we have validated TRE-based viral effectors of oChiEF (ChR2 variant) ^28^, NpHR ^29^, GCaMP6s ^30^ and HTG (histoneGFP-TVA-Glycoprotein) ^26^, which can be broadly utilized in various conditions with viral serotypes, promoters and transgenic animals. As the design of splitting tTA is also applicable for splitting rtTA (Fig. S1B-F), there could be more adaptable options for intersectional control in various experimental purposes.

The intein mediated splicing technique has long been used before, but the efficiency is varying depending on the intein sequence and the application context. We adopted a so-far best strategy based on Gp41-1 sequences because they have small sizes and possess the most rapid reaction rate among all split inteins examined ^17, 20^. In the future, mapping split site of Gp41-1 amino acid sequence for more efficient splicing and compatible segmentation of various controllers might bring further improved performance for IBIST tools. We have validated the specificity of IBIST tools by careful evaluation and noticed a small bit of leak by AAV-tTAN or AAV-tTAC when the titer was higher than 10^13^. While care should be taken when using all AAV vectors at high titers, we recommend to simply use AAVs with titers at the level of 10^12^ in order to avoid potential non-specific leak. Future work may further ensure the specificity by fusing tTAN and tTAC with destabilizing domains such as ddFKBP and SopE ^38-41^ which help to clear residual of tTAN or tTAC in the cells accordingly. In our experiences, some TRE effectors do have faint baseline expression of fluorophore tag and hence are only suitable for functional studies with high-level expression demand. Therefore, the strength of our system lies more in the specific and sufficient gene expression for effectors such as optogenetic opsins and Ca^2+^ indicators.

Our system is not only compatible with increasing mouse driver lines with Cre and Flp for most major cell types in the brain ^5, 6, 23^, but also complementary for the existing intersectional tools such as Cre- and Flp-dependent constructs created by pioneering works ^4, 11^. Intersectional control with five or more features (Fig. S7) becomes possible if the split tTA fragments are built upon those Cre- and Flp-dependent constructs ^11^, antero- and retrograde tracing or other promoter and enhancer dependent constructs ^42, 43^. Intersectional ON and OFF is also possible by the IBIST tools if one of the tTA fragments is flanked by palindromic sites recognized by site-specific recombinases (e.g. Cre-loxp site-specific recombination system). As the intein-split strategy could be adapted for other controllers such as Cre ^7^, it would be possible to devise two orthogonal strategies in the same animal by independent and parallel synthesis of two single controllers in intersectional manners, respectively. Given that the IBIST tools can be entirely implemented in viral vectors including AAV and rabies vectors, our system offers experimental conveniences and also holds great promise as intersectional tools for animal species with limited transgenic resources such as primates.

## Acknowledgments

We thank all members of the Xu lab and the Long Lab. We thank non-human primate facility for providing cynomolgus monkey, imaging facility and animal facility for technical support, B. Roska, Z.-K. Qian, H.-K. Zeng, M.-M. Luo and L.-Q. Luo for sharing plasmids, E.M. Callaway for sharing rabies-related cell lines and plasmids. This study was supported by the Strategic Priority Research Program of the Chinese Academy of Sciences (XDB32010100), Shanghai Municipal Science and Technology Major Project (2018SHZDZX05), the National Natural Science Foundation of China (31771180 and 91732106) and the international collaborative project (19490711800) of Shanghai Science and Technology Committee.

---

**Supplementary Figure 1.**
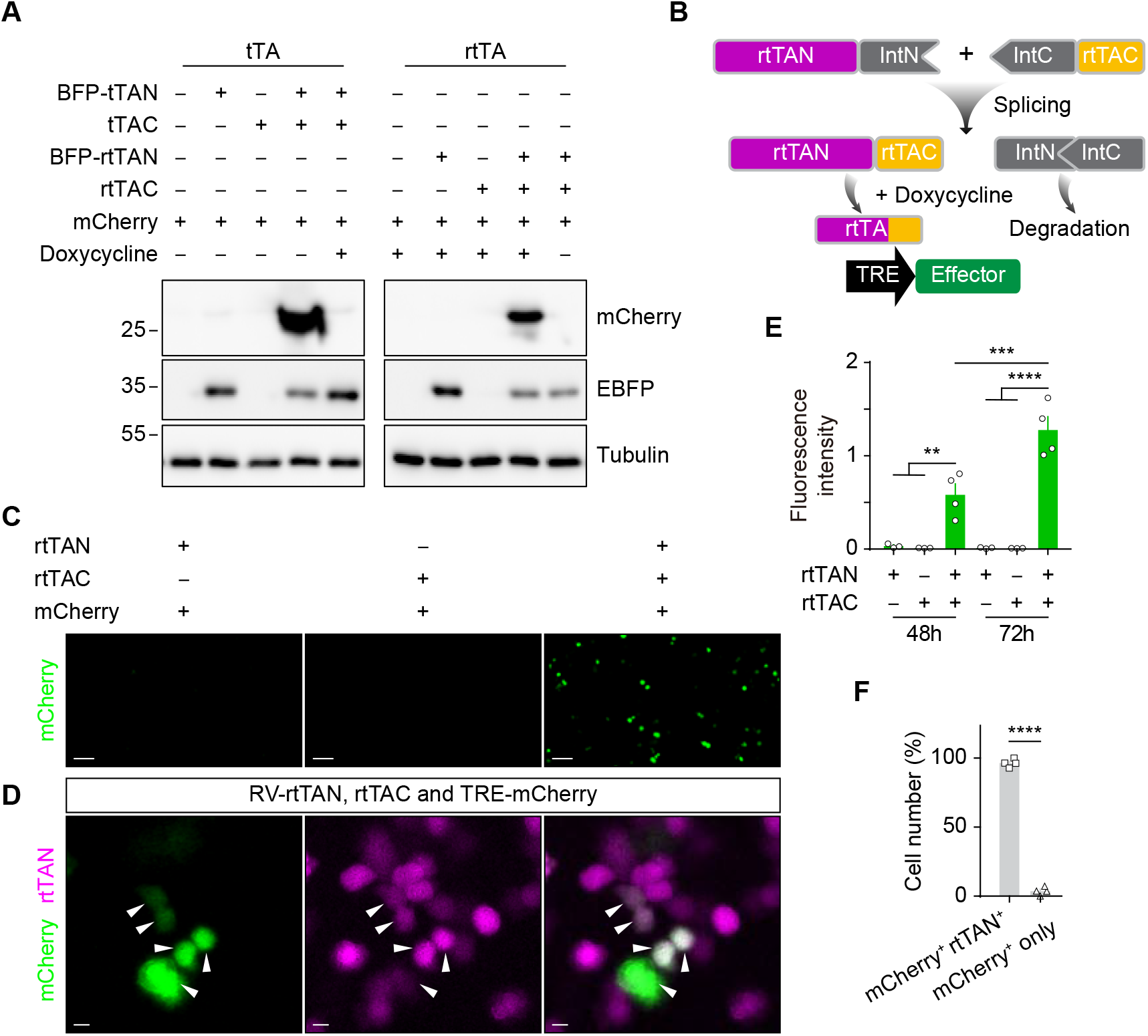
The specific reconstitution of rtTA *in vitro.* **A**, Western blot analysis of mCherry and BFP tagged tTA or rtTA after plasmids transfection in HEK293T cells. **B**, Diagram depicting the reconstitution of rtTA by intein-based protein splicing. **C**, Examples of mCherry fluorescence (shown in green) in HEK293T cells 72 h after transfection with reporter (TRE-mCherry) and rtTA plasmids and infection with or without RV-rtTAN (GFP tagged). Scale bars: 100 μm. **D**, Examples showing the fluorescence of TRE-mCherry (shown in green) and RV-rtTAN (shown in magenta) in HEK293T cells with all plasmids transfected. Arrows: co-labeled cells. Scale bars: 10 μm. **E**, Summary of red fluorescence intensity of HEK293T cells at 48 h and 72 h post transfection in **C** (RV-rtTAN only, n = 3 FOV; rtTAC only, n = 3 FOV; rtTAC+RV-rtTAN, n = 4 FOV). One-way ANOVA revealed a significant difference between groups (F_(5,14)_ = 34.39, P < 0.0001). Tukey's multiple comparisons test revealed that fluorescence intensity in rtTAC+RV-rtTAN group at 72 h is significantly higher than that at 48 h post transfection (P < 0.001), and both are significantly higher than other groups at the same time post transfection (P < 0.01 for 48 h and P < 0.0001 for 72 h). **F,** Percentage summary of cells labeled by TRE-mCherry and RV-rtTAN in **D** (96.4±1.5% co-labeled vs. 3.6±1.5% reporter only; paired *t*-test, P < 0.0001, n = 4 FOV).

**Supplementary Figure 2.**
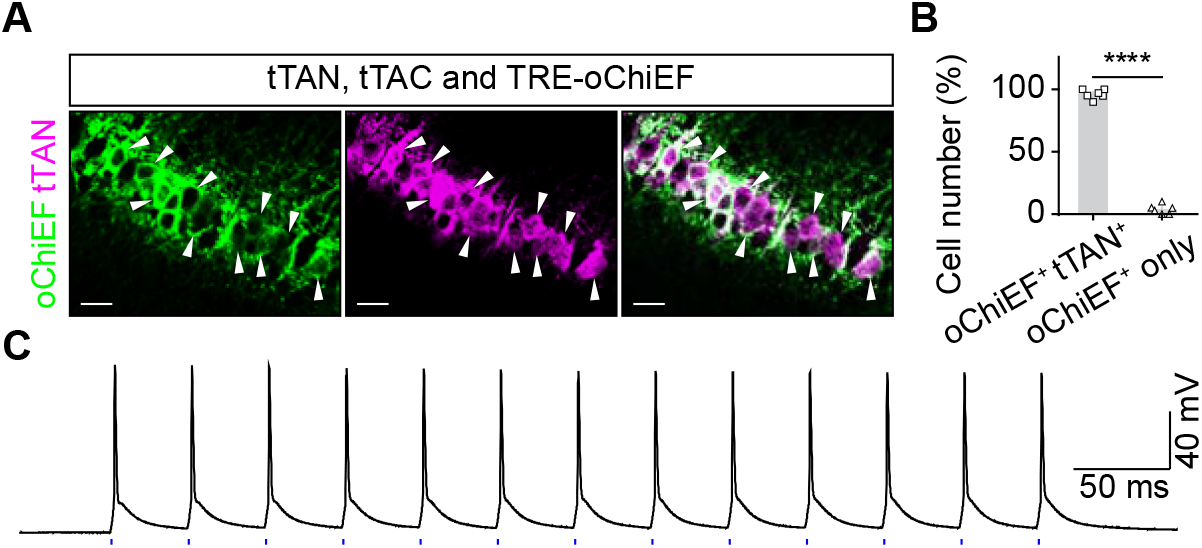
The histology of IBIST-based optogenetic opsins. **A**, Confocal images showing fluorescent labeling by AAV-CaMKIIα-tTAN-BFP (shown in magenta) and AAV-TRE-oChiEF-mCherry (shown in green) in hippocampal sections from animal injected with all three AAVs. Scale bars: 20 μm. **B**, Percentage summary of cells labeled by AAV-TRE-oChiEF and AAV-CaMKIIα-tTAN-BFP (96.2±1.6% co-labeled vs. 3.8±1.6% reporter only; paired *t*-test, P < 0.0001, n = 6 FOV, N = 2 animals). **C**, Example showing light-evoked spikes in hippocampal cells infected with AAV-TRE-oChiEF and AAVs (CaMKIIα promoter) of tTAN and tTAC. Blue ticks indicate blue LED light pulses (3 mW, 2 ms) at 25 Hz. Similar recordings were replicated in 3 cells.

**Supplementary Figure 3.**
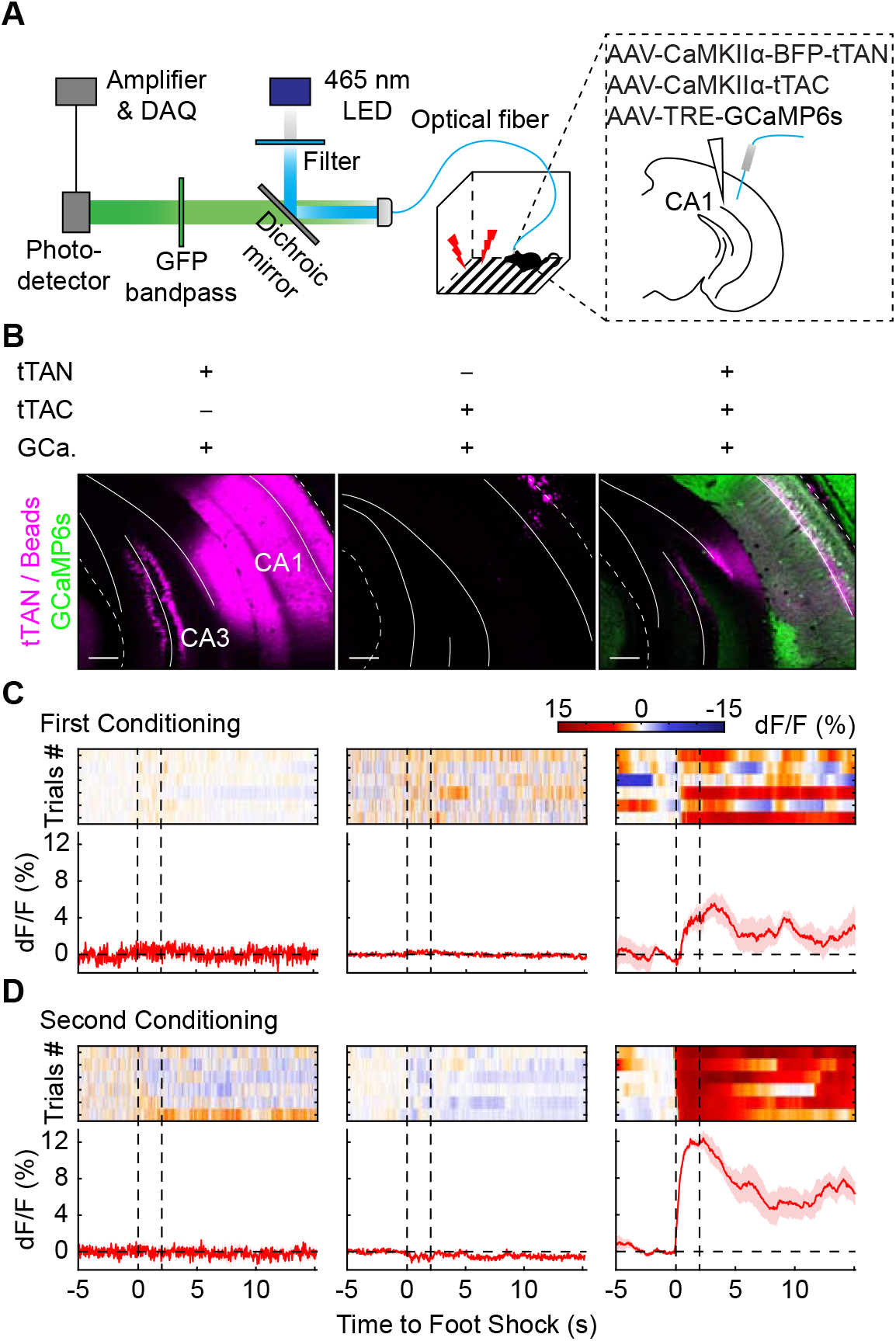
The IBIST-based Ca^2+^ recording by GCamP6s. **A**, Scheme depicting the photometric recording in vCA1 injected with AAVs of TRE-GCaMP6s, CaMKIIα-tTAN and CaMKIIα-tTAC. **B**, Examples showing fluorescence of hippocampal slice from animal injected with part or all AAVs of tTAN (tagged with BFP, shown in magenta), tTAC and TRE-GCaMP6s (shown in green) and co-injected with blue beads (shown in magenta). Scale bars: 200 μm. **C and D**, Examples of heatmaps and averaged traces of Ca^2+^ signals upon foot shocks (between vertical dash lines, 6 trials) in the first (**C**) and second (**D**) fear conditioning session recorded from animal injected in **B**. Similar histological and photometric recording results were obtained from 1 – 2 replicates in each group (tTAC, N= 1 animal; tTAN, N= 2 animals; tTAC+tTAN, N= 2 animals).

**Supplementary Figure 4.**
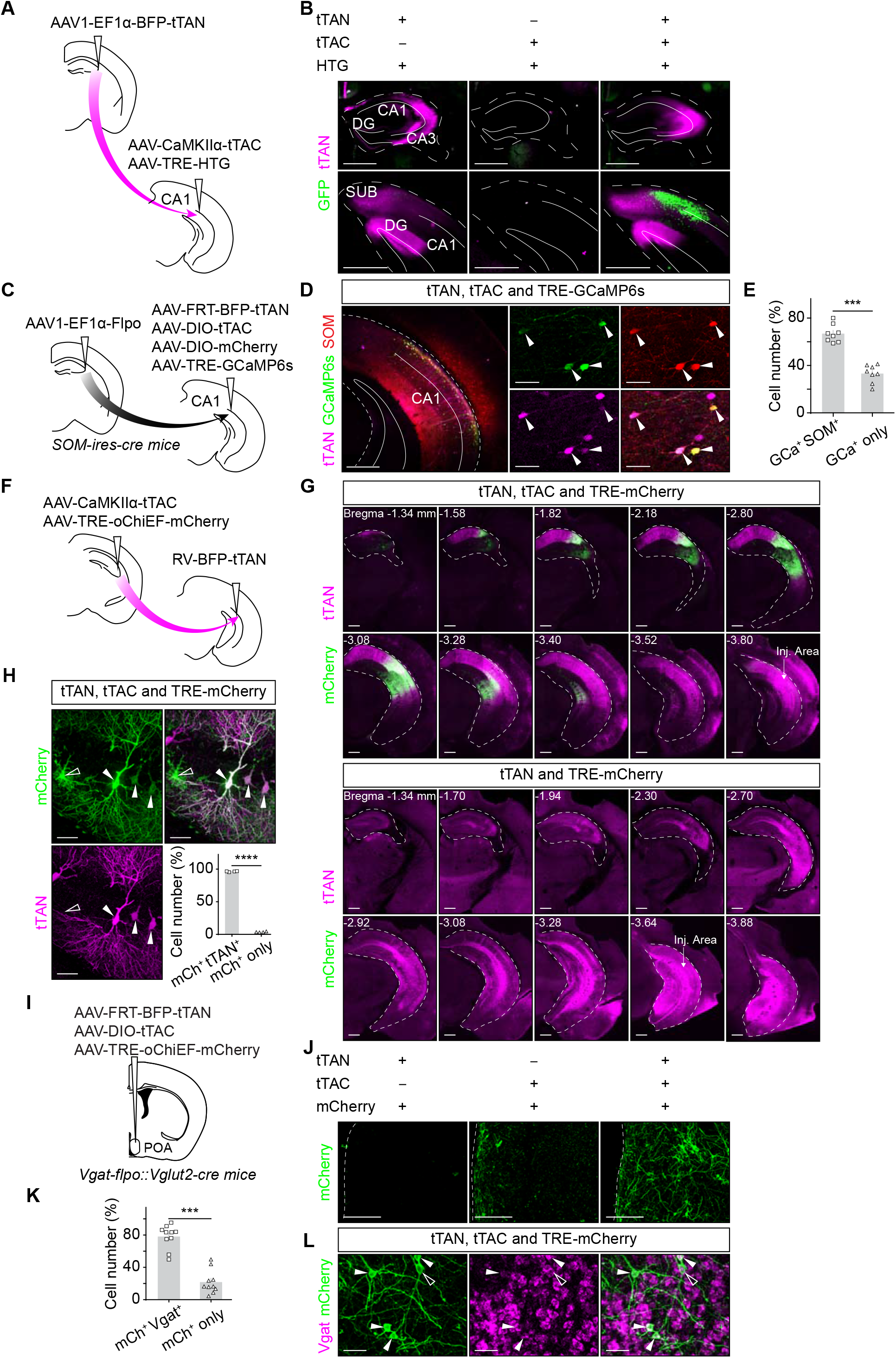
The IBIST-based fluorophore labeling by two features. **A,** Scheme illustrating AAV injections to define cells by two features: anterograde transsynaptic tracing and pyramidal-cell-specific fluorophore labeling. **B**, Examples of fluorescent labeling from animal injected in **A**. Scale bars: 200 μm. **C,** Scheme illustrating AAV injections to combine anterograde transsynaptic tracing with SOM+ interneuron-specific fluorophore labeling. **D**, *Left*, examples showing fluorescence in ventral hippocampus from *SOM-ires-cre* animal injected with all the AAVs. Scale bars: 500 μm. *Right*, enlarged pictures showing co-labeling by SOM positive cells (shown in red) and AAV-TRE-GCaMP6s (shown in green). Filled arrow, double labeled. Scale bars: 50 μm. **E,** Percentage summary of cells labeled by AAV-DIO-mCherry and AAV-TRE-GCaMP6s (66.9±2.7% co-labeled vs. 33.1±2.7% reporter only; paired *t*-test, P = 0.0004, n = 8 FOV, N = 2 animals). **F,** Scheme illustrating AAV injections to combine retrograde transsynaptic rabies tracing with pyramidal-cell-specific fluorophore labeling. **G**, *Top*, examples showing dorsal and ventral hippocampus cells labeled by rabies-BFP-tTAN (shown in magenta) and AAV-TRE-oChiEF-mCherry (shown in green). *Bottom*, examples showing retrograde labeling in the absence of AAV-CaMKIIα-tTAC as control. The arrows indicate injection area. Scale bars: 500 μm. **H**, Examples and summary showing fluorescent labeling from animals in **G** (top). Open arrow, reporter only. Filled arrow, double labeled. Scale bars: 50 μm. 96.3±0.4% of mCherry+ cells were co-labeled by rabies-BFP-tTAN and 3.7±0.4% were not co-labeled (paired *t*-test, P < 0.0001, n = 4 FOV). **I**, Scheme illustrating AAV injections into the hypothalamic preoptic area (POA). **J**, Examples showing mCherry fluorescence (shown in green) in POA injected with AAV-TRE-oChiEF and either AAVs of AAV-FRT-tTAN, AAV-DIO-tTAC or both. Scale bars: 200 μm. **K**, Percentage summary of cells labeled by mCherry and *Vgat* (78.0±4.6% co-labeled vs. 22.0±4.6% reporter only; paired *t*-test, P = 0.0002, n = 10 FOV). **L**, Examples showing in situ hybridization of *Vgat* and immunostaining of mCherry in animal with all AAVs injected. Open arrow, reporter only. Filled arrow, double labeled. Scale bars: 50 μm.

**Supplementary Figure 5.**
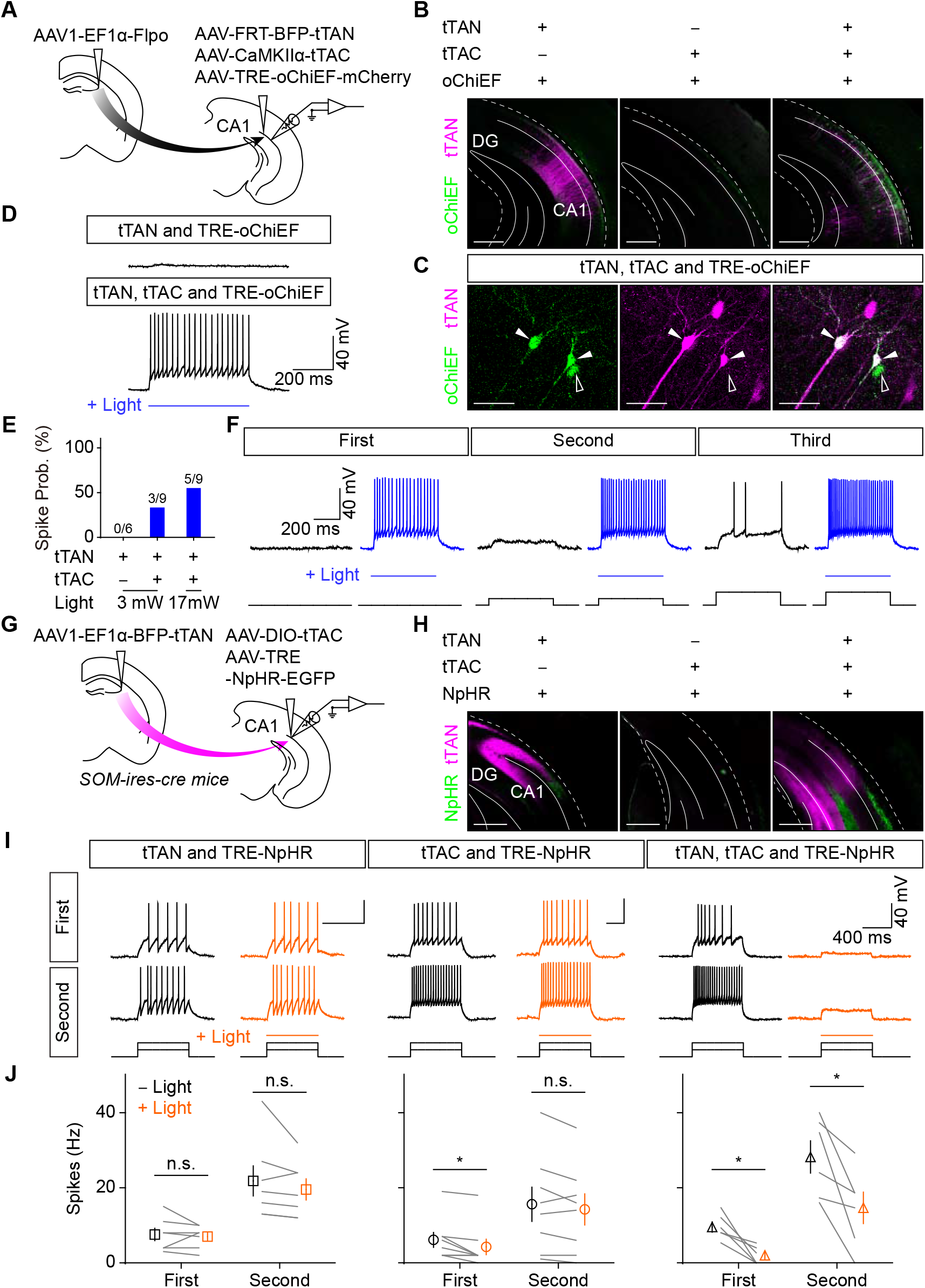
The IBIST-based optogenetic manipulation for cells defined by two features. **A**, Scheme illustrating AAV injections to define cells by two features: anterograde tracing and pyramidal-cell-specific expression of ChR2. **B**, Examples of blue and red fluorescence in ventral hippocampus from animals injected with AAVs of tTAN (shown in magenta), tTAC or TRE-oChiEF-mCherry (shown in green). Scale bars: 500 μm. **C**, Examples showing whole-cell current-clamp recording in brain slices with blue LED light stimulation (blue line). All groups were injected with AAV-TRE-oChiEF and co-injected with AAVs of either tTAN, tTAC, or both (also see Fig. 2f, tTAC and TRE-oChiEF). **D**, Examples showing fluorescent labeling by AAV-FRT-BFP-tTAN (shown in magenta) and AAV-TRE-oChiEF-mCherry (shown in green) in hippocampal sections from animal injected with all three AAVs. Open arrow, reporter only. Filled arrow, double labeled. Scale bars: 50 μm. **E,** Summary of spike probabilities (prob.) in hippocampal cells evoked by whole-field blue LED light from animals injected with all the AAVs or with tTAC omission: tTAN, N = 2 animals; tTAN+tTAC, N = 1 animal. **F**, Examples showing whole-cell current-clamp recording from hippocampal cells in brain slices upon current injections (0, 40 and 80 pA; blue with light; black without light). **G**, Scheme illustrating AAV injections to combine anterograde tracing with SOM+ interneuron specific expression of NpHR in *SOM-ires-Cre* mice. **H**, Examples of blue and green fluorescence in dorsal and ventral hippocampus from animal injected with AAVs of tTAN (shown in magenta), tTAC or TRE-NpHR-EGFP (shown in green). Scale bars: 500 μm. **I**, Examples showing whole-cell current-clamp recording from hippocampal cells in brain slices upon current injections with 40 pA steps (first and second sweeps with prominent spikes; yellow with light; black without light). **J,** Spike frequency in hippocampal cells in the absence (black) or presence (yellow) of 589 nm laser light (18 mW) from animals injected with all the AAVs (Wilcoxon matched-paired signed rank test; first sweep, 9.6±1.4 Hz vs. 2.0±1.0 Hz with light; P = 0.0313; second sweep, 28.2±4.3 Hz vs. 14.7±4.2 Hz with light; P = 0.0313, N = 2 animals). The tTAN animals: Wilcoxon matched-paired signed rank test; first sweep, 7.6±1.6 Hz vs. 7.0±1.1 Hz with light; P = 0.5; second sweep, 21.9±4.0 Hz vs. 19.6±2.8 Hz with light; P = 0.1875, N = 2 animals. The tTAC animals: Wilcoxon matched-paired signed rank test; first sweep, 6.1±2.0 Hz vs. 4.3±2.1 Hz with light; P = 0.0313; second sweep, 15.6±4.6 Hz vs. 14.3±4.2 Hz with light; P = 0.1719, N = 2 animals.

**Supplementary Figure 6.**
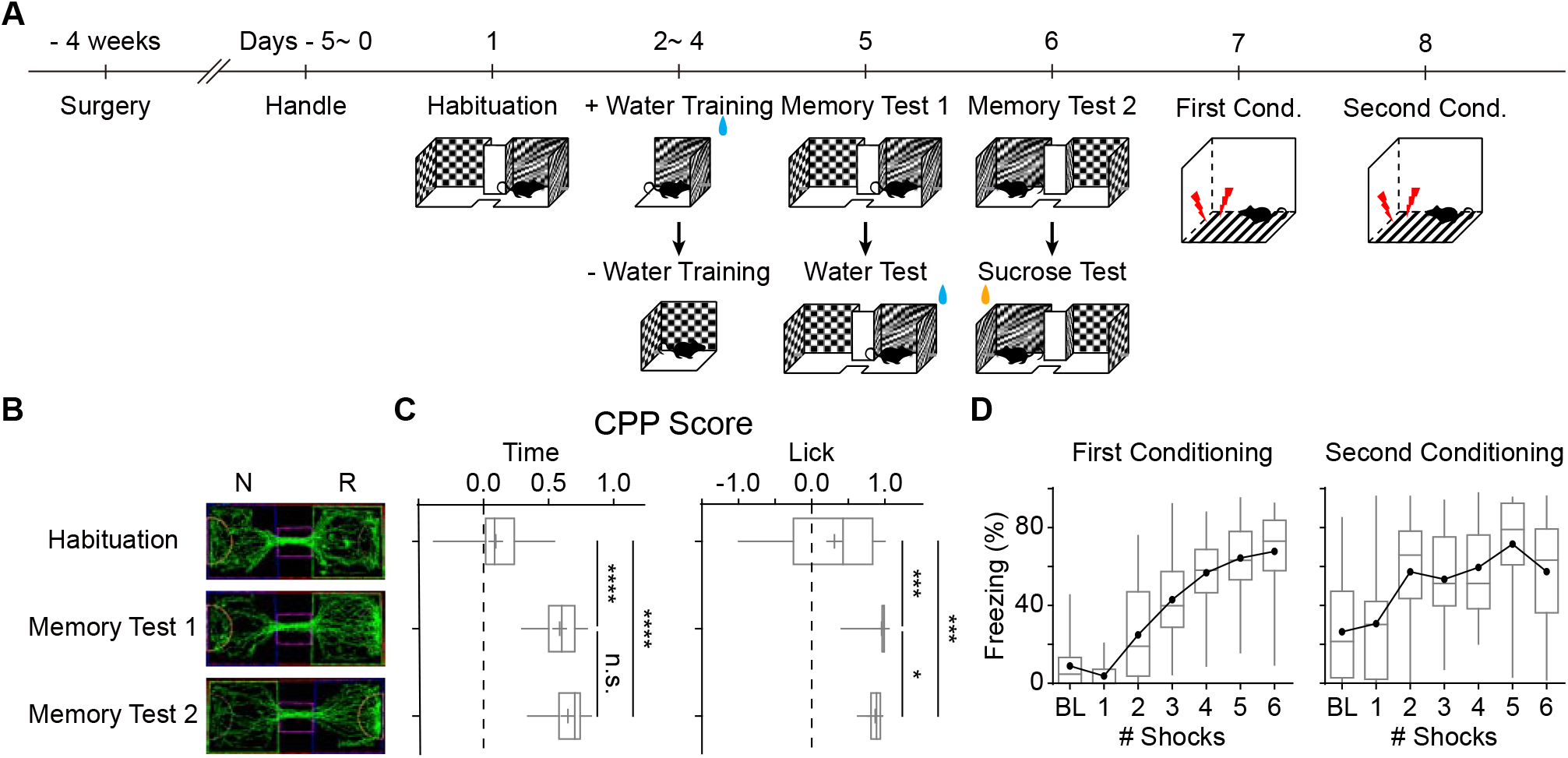
Behavioral summary of CPP and fear conditioning. **A**, Scheme illustrating behavioral training protocols for CPP and fear conditioning. **B**, Examples showing trajectories during test sessions of water CPP. N, non-rewarding context. R, rewarding context. **C**, Summary of CPP score during test sessions. One-way ANOVA revealed a significant difference in CPP scores after training (time: F_(1.868, 41.09)_ = 78.25, P < 0.0001; licks: F_(1.1, 24.2)_ = 22.37, P < 0.0001). **D**, Percentage of freezing time during first conditioning and second conditioning. BL, baseline.

**Supplementary Figure 7.**
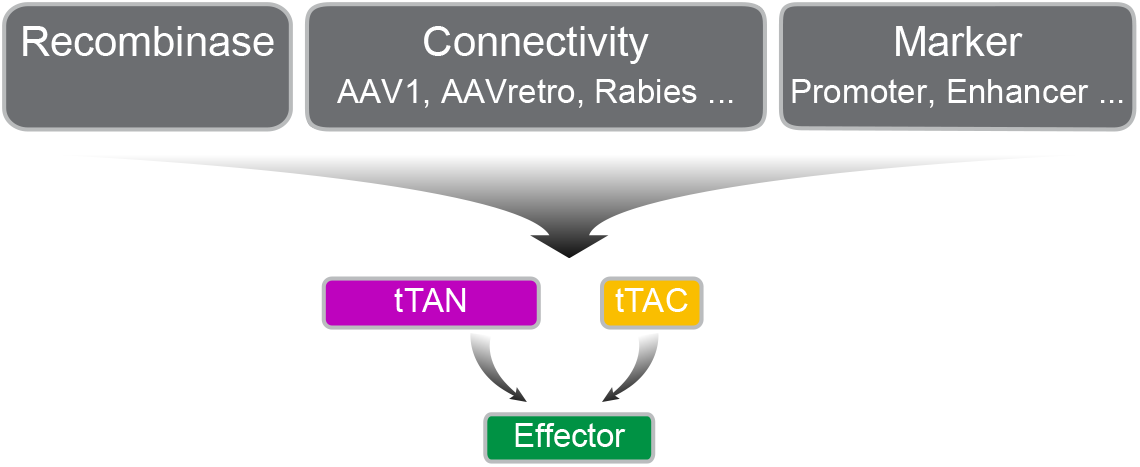
Multiple-feature strategy by IBIST. Scheme illustrating strategies for IBIST-based labeling cells via multiple features. There are two layers of controllers: one consists of intein-split tTA fragments; the other are recombinase, viral tracer and molecular markers. Similar design is also applicable for intein-split rtTA fragments. Both tTA and rtTA offer an additional feature to be controlled by doxycycline.

## Author contributions

H.-S.C., X.-L.Z., R.-R.Y., G.L. and C.X. conceived the project. G.L. and C.X. supervised the work in the study. X.-L.Z. designed and tested DNA constructs of split controller, performed Western blot and analyzed the results. R.-R.Y. generated the rabies vectors, performed rabies-related experiments and analyzed the results. H.-S.C., G.-L.W., R.-R.Y. N.Z. and X.-Y.Z. injected the AAV vectors, performed the histology and confocal imaging and analyzed the results. G.-L.W. performed the electrophysiological recording from acute brain slices and analyzed the results. H.-S.C. and S.Q. performed the photometric recording and analyzed the results. Y.-F.X. and Z.-M.S. performed the surgery in the primate. D.-Y.W. and X.-H.X. performed AAV injection and in situ hybridization and analyzed the results. H.-S.C., G.L. and C.X. wrote the manuscript with inputs from all authors.

## Methods

### Plasmids

All vectors in this study were constructed using EasyGeno Assembly Cloning kit (TIANGEN, Beijing) according to the manual, except for the specially noted plasmids. Briefly, the backbone plasmid was linearized and the insert fragments was amplified through normal polymerase chain reactions. All DNA products were separated in agarose gel to confirm sizes and following purified using TIANgel Midi Purification Kit (TIANGEN). The mixture of 0.01 pM of backbone and 0.03 pM of each insert fragments were delivered for transformation in Stable3 competent bacteria after 30 minutes incubation in 50 °C. All the plasmids are verified by sequencing.

The plasmids created in this study will be available on Addgene. The response plasmids for Tet-On and Tet-Off gene expression all contain Tet-responsive Ptight promoter. The source of backbone plasmids and template plasmids for created plasmids were summarized in the supplementary Excel file.

In this study, pAAV-TRE-HTG (Addgene, #27437) was a gift from L. Luo ^26^; pAAV-EF1α-DIO-TVA950-T2A-CVS11G was a gift from B. Roska ^44^; pcDNA-B19N, pcDNA-B19P, pcDNA-B19L, pcDNA-B19G and pSADdeltaG-F3 (#32634) were gifts from E.M. Callaway ^45^; pLenti-CaMKIIα-VChR1-EYFP (#20954) was a gift from K. Deisseroth ^46^; pLKO-GFP-IntN-SopE and pLKO-IntC-Flag were gifts from Z.-K. Qian ^47^; pmSyn1-EBFP-Cre (#51507) was a gift from H. Zeng ^6^; pAAV-TRE-fDIO-GFP-IRES-tTA (#118026) was a gift from M. Luo ^48^.

### Virus preparations

#### Rabies vectors

The SADΔG rabies virus was generated as described before ^34^. In brief, G-deleted rabies virus expressing rtTAN-GFP (RV-rtTAN) was recovered by transfecting B7GG cells (kindly provided by Ed Callaway, Salk Institute) with pcDNA-B19N, pcDNA-B19P, pcDNA-B19L, pcDNA-B19G and pRabiesΔG-rtTAN-GFP in a humidified atmosphere of 5% CO_2_ and 95% air at 35°C. The recovered RV-rtTAN was amplified in B7GG cells, the supernatant was used to infect HEK293T cells. The titer of the RV-rtTAN viral supernatant was in the range of 10^3^ – 10^4^ infectious units/mL.

The G-deleted rabies virus expressing tTAN-BFP (RV-tTAN) was recovered similarly. For *in vivo* injection, RV-tTAN containing supernatant was concentrated by 25,000 rpm centrifuge for 4 h. The titer was in the range 10^6^ – 10^7^ infectious units/mL.

#### AAV vectors

AAV vectors were typically generated with serotype 8 and 9, unless otherwise noted in the text. In this study, the effector AAVs for Tet-On and Tet-Off gene expression all contain Tet-responsive Ptight promoter.

Most AAV vectors were produced by the gene editing core facility at the Institute of Neuroscience: AAV2/9-CaMKIIα-BFP-tTAN (7.2×10^13^), AAV2/9-CaMKIIα-tTAC (4.2×10^13^), AAV2/1-EF1α-BFP-tTAN (4.3×10^13^), AAV2/1-EF1α-Flpo (8×10^12^), AAV2/8-EF1α-FRT-BFP-tTAN (8.9×10^13^), AAV2/8-EF1α-DIO-tTAC (1.4×10^14^), AAV2/8-EF1α-DIO-mCherry (2.4×10^13^), AAVretro-CaMKIIα-BFP-tTAN (1.8×10^13^), AAVretro-CaMKIIα-tTAC (6.8×10^12^), AAV2/9-TRE-HTG (1.8×10^13^), AAVretro-EF1α-Flpo (7.9×10^12^), AAVretro-EF1α-FRT-BFP-tTAN (1.3×10^13^), AAVretro-EF1α-DIO-tTAC (7.5×10^12^), AAVretro-CaMKIIα-cre (2.4×10^13^). The TRE-driven AAV effectors were produced with serotype 9 (Taitool Bioscience, Shanghai): AAV2/9-TRE-oChiEF-mCherry (1.59×10^13^), AAV2/9-TRE-GCaMP6s (2×10^13^), AAV2/9-TRE-NpHR-EGFP (1,5×10^13^). The titer of AAVs was typically diluted into the range of 10^12^ before *in vivo* injection.

### Cell culture

HEK293T and BG77 cells (provided by E.M. Callaway) are cultured with 10% (v/v) FBS/DMEM at 5% CO_2_ and 37 °C.

#### Reconstitution of tTA in vitro

The HEK293T cells were divided into four groups and transfected with plasmids by Lipofectamine 2000 (11668-027, Invitrogen). The four groups (tTAN, tTAC, both or neither) of plasmids for transfection consist of reporter pAAV-TRE-HTG and intein-split tTA plasmids with EF1α promoter. The fluorescence images for HEK293T cells were acquired at 48 h and 72 h after the transfection.

#### Reconstitution of rtTA *in vitro*

The HEK293T cells were divided into three groups and transfected with plasmids of pAAV-TRE-HTG and pAAV-EF1α-rtTAC by Lipofectamine 2000 (11668-027, Invitrogen). The culture medium was replaced with doxycycline-containing medium (100 ng/ml, MedChemExpress, HY-N0565B) 6 h after the transfection. Eighteen hours later, the culture medium was replaced with RV-rtTAN-containing medium (rabies-containing supernatant from B7GG cells) for one group of cells and incubated for 6 h, and then replaced with fresh medium again (with doxycycline 100 ng/ml). For the control groups, pAAV-EF1α-rtTAC plasmid or RV-rtTAN-containing medium was omitted, respectively. The fluorescence images for HEK293T cells were acquired at 48 h and 72 h after the transfection.

#### Quantification for fluorescence and co-labeling *in vitro*

The fluorescence images for HEK293T cells were acquired by Olympus microscope (CKX53 with X-CITE LED, 10X objective, same exposure settings for the same type of experiment). One FOV image per culture plate was taken for quantification (green for TRE-HTG and rtTAN-GFP, red for TRE-mCherry and blue for tTAN-BFP). The mean fluorescence intensity in the whole FOV was measured in ImageJ. The reporter cells and co-labeled cells were counted in Imaris (v7.3, Bitplane).

### Animals

Animals were housed under a 12 h light / dark cycle and provided with food and water *ad libitum* in the Institute of Neuroscience animal facility (mice and cynomolgus monkey). All animal procedures were performed in accordance with institutional guidelines and were approved by the Institutional Animal Care and Use Committee (IACUC) of the Institute of Neuroscience (CAS Center for Excellence in Brain Science and Intelligence Technology), Chinese Academy of Sciences. Wild-type C57BL/6J (Slac Laboratory Animal, Shanghai), Slc32a1^tm1.1(flpo)Hze^ (Vgat^flpo^, #029591), Slc17a6^tm2(cre)Lowl/J^(Vglut2^cre^, #016963) and *SOM-ires-Cre*^*13*^ mice were used. The Vglut2-cre/Vgat-flpo mouse line was obtained by crossing Vglut2-cre and Vgat-flpo animals. Mice used in the experiment were heterozygous. All of the experimental mice used in the study were adult male mice (over 8 weeks). One male cynomolgus monkey (*Macaca fascicularis*, 4.2 kg, 13 years old) was obtained from non-human primate facility of Institute of Neuroscience after its full use in reproductive research.

### Stereotaxic injections

Mice were anaesthetized with isoflurane (induction 5%, maintenance 2%, RWD R510IP, China) and fixed in a stereotactic frame (RWD, China). Before the surgery, local analgesics (Lidocaine, H37022147, Shandong Hualu Pharmaceutical) were administered. A feedback-controlled heating pad (FHC) was used to maintain the body temperature at 35°C. Glass pipettes (tip diameter 10 – 20 μm) connected to a Picospritzer III (Parker Hannifin Corporation) were filled with virus solutions and injected at following coordinates (posterior to Bregma, AP; lateral to the midline, LAT; below the brain surface, DV; in mm): mPFC: AP +1.98, LAT ±0.6, DV −1.62, angle 10°; Amygdala: AP −1.2, LAT ± 3.0, DV −4.35; NAc: AP +1.0, LAT ± 1.0, DV −4.2; Hypothalamic POA: AP +0.35, LAT +0.3, DV −5.1; dorsal CA3: AP −1.5, LAT ± 2.1, DV −1.7; ventral CA1: AP −3.2, LAT ± 3.4, DV −1.5 or 3.5. To identify the injection site, the virus solution was typically mixed at 1:500 with blue beads (fluorescent polymer microspheres, Thermo Scientific).

#### Local injections in hippocampus

A mixture (300 nL) of AAV2/9-TRE-HTG, AAV2/9-CaMKIIα-tTAC and AAV2/9-CaMKIIα-tTAN-BFP (ratio 1:1:1) were injected into dorsal CA3 (AP - 1.5, LAT ± 2.1, DV −1.7). In control groups, one of these three viruses (AAV2/9-TRE-HTG, AAV2/9-CaMKIIα-tTAC and AAV2/9-CaMKIIα-tTAN-BFP) was omitted for injections (ratio 1:1 for two AAVs). In negative control group, only AAV2/9-TRE-HTG (300 nL) was injected into dorsal CA3.

For electrophysiological experiments combined with optogenetics, same AAVs were injected into ventral CA1 (AP −3.2, LAT ±3.4, DV −1.5), except that AAV2/9-TRE-oChiEF-mCherry or AAV2/9-TRE-NpHR-EGFP was used instead of AAV2/9-TRE-HTG.

For photometric experiments, same AAVs were injected into ventral CA1 (AP −3.2, LAT ±3.4, DV −1.5), except that AAV2/9-TRE-GCaMP6s was used instead of AAV2/9-TRE-HTG.

#### Local injections in hypothalamus

A mixture (300 nL) of AAV2/9-TRE-oChiEF-mCherry, AAV2/8-DIO-tTAC and AAV2/8-FRT-tTAN (ratio 1:1:1) were injected into POA (AP +0.35, LAT ± 0.5, DV −5.1). In control groups, one of these three viruses was omitted for injections (ratio 1:1 for two AAVs).

#### Retrograde tracing from ventral hippocampus to dCA3

A Mixture (300 nL) of AAV2/9-CaMKIIα-tTAC and AAV2/9-TRE-oChiEF-mCherry (ratio 1:1) were injected into dCA3 (AP −1.5, LAT ± 2.1, DV −1.7), while RV-BFP-tTAN (500 nL) was injected into vCA1 (AP −3.2, LAT ±3.4, DV −3.5) simultaneously. In control group, tTAC was omitted for injection.

#### Retrograde tracing from downstreams of ventral hippocampus

To define hippocampal cells by two projection targets and CaMKIIα promotor, AAVretro-CaMKIIα-tTAC (200 nL), AAVretro-CaMKIIα-BFP-tTAN (200 nL) were injected into two output regions or co-injected at the following coordinates: mPFC (AP +1.98, LAT ±0.6, DV −1.62, angle 10°)s, NAc (AP +1.0, LAT ±1.0, DV −4.2), and amygdala (AP −1.2, LAT ±3.0, DV −4.35). AAV2/9-TRE-GCaMP6s (200 nL) were injected into ventral CA1 (AP −3.2, LAT ±3.4, DV −3.5). In control groups, either tTAC or tTAN was omitted for injection. All AAVs were co-injected with blue beads.

To define hippocampal cells by four projection targets and CaMKIIα promotor, AAVretro-EF1α-Flpo (200 nL), AAVretro-EF1α-FRT-BFP-tTAN (200 nL), AAVretro-CaMKIIα-cre (200 nL), and AAVretro-EF1α-DIO-tTAC (200 nL) were injected into four regions at the following coordinates respectively: mPFC (AP +1.98, LAT ±0.6, DV −1.62, angle 10°), NAc (AP +1.0, LAT ±1.0, DV −4.2), amygdala (AP −1.2, LAT ±3.0, DV −4.35), and lateral septum (AP +0.5, LAT ±0.35, DV −2.75). All AAVs were co-injected with blue beads.

#### Anterograde tracing from dorsal CA3 to ventral hippocampus

AAV2/1-EF1α-BFP-tTAN (300 nL) were injected into dorsal CA3 (AP −1.5, LAT ± 2.1, DV −1.7). Then a mixture (400 nL) of AAV2/9-TRE-HTG and AAV2/9-CaMKIIα-tTAC (ratio 1:1) were injected into ventral CA1 (AP −3.2, LAT ±3.4, DV −1.5). For controls, either tTAC or tTAN was omitted for injection.

To label SOM+ interneuron with dCA3 inputs, AAV2/1-EF1α-Flpo and AAV2/9-TRE-GCaMP6s were used instead of AAV2/1-EF1α-BFP-tTAN and AAV2/9-TRE-HTG, respectively. N terminal of tTA was expressed in a Flpo-dependent manner by injecting AAV2/8-EF1α-FRT-BFP-tTAN (300 nL) into ventral CA1 (AP −3.2, LAT ±3.4, DV −1.5). A mixture (300 nL) of AAV2/8-EF1α-DIO-tTAC and AAV2/8-EF1α-DIO-mCherry (ratio 1:1) were delivered in ventral CA1 to label SOM+ cells.

For photometric experiments, same AAVs were injected into dorsal CA3 (AP −1.5, LAT ± 2.1, DV −1.7) and ventral CA1 (AP −3.2, LAT ±3.4, DV −1.5) as fluorophore labelling in SOM+ interneuron, except that AAV2/9-CaMKIIα-tTAC was used instead of the mixture of AAV2/8-EF1α-DIO-tTAC and AAV2/8-EF1α-DIO-mCherry.

For electrophysiological experiments combined with optogenetics excitation, same AAVs were injected into dorsal CA3 (AP −1.5, LAT ± 2.1, DV −1.7) and ventral CA1 (AP −3.2, LAT ±3.4, DV −1.5) as photometric experiments, except that AAV2/9-TRE-oChiEF-mCherry was used instead of AAV2/9-TRE-GCaMP6s.

For electrophysiological experiments combined with optogenetics inhibition in SOM+ interneuron, AAV2/1-EF1α-BFP-tTAN (300 nL) were injected into dorsal CA3 (AP −1.5, LAT ± 2.1, DV −1.7). Then a mixture (400 nL) of AAV2/9-TRE-NpHR-EGFP and AAV2/8-EF1α-DIO-tTAC (ratio 1:1) were injected into ventral CA1 (AP −3.2, LAT ±3.4, DV −1.5).

#### Local injections in primate cortex

One cynomolgus monkey was under general anesthesia (isoflurane 1.5~3%) and sterile conditions after subcutaneous injection of atropine sulfate (0.1mg /kg), intra-muscular injection of tiletamine hydrochloride and zolazepam hydrochloride (0.1mg/kg) and tracheal intubation. The electrocardiogram (ECG), pulse oximeter (SpO2) and end-tidal carbon dioxide (CO_2_) were monitored continuously. The core temperature was maintained at around 38°C by an electric blanket. A craniotomy (1.5×2.0 cm^2^) was opened in the skull over primary visual cortex. The dura was opened to expose the cortex at 4 sites (~5 mm in between). The virus solutions in glass pipettes were injected (30 nL/min) into the cortex at a depth of ~1.5 mm. A mixture (600 nL) of AAV2/9-TRE-oChiEF-mCherry, AAV2/9-CaMKIIα-tTAC and AAV2/9-CaMKIIα-tTAN-BFP (ratio 1:1:1) were injected into two sites. In the other two sites, either AAV2/9-CaMKIIα-tTAC or AAV2/9-CaMKIIα-tTAN-BFP was omitted for injections. The craniotomy was stabilized by sterile plastic sheets during injection, and then was covered with tissue glue and acrylic cement. Animal was treated with antibiotics and analgesia and returned to its home cage after the surgery. Slice recording was performed 6 weeks after the AAV injection.

### Histology and quantification for fluorescence and co-labeling

One to four weeks after viral injections, mice were transcardially perfused with phosphate buffered saline (PBS) followed by 4% (w/v) paraformaldehyde (PFA) in PBS. Brains were post-fixed in PFA overnight at 4°C. and then cut into 80 μm thick coronal slices with a vibratome (Leica, Germany).

For immunostaining of free-floating sections, sections were incubated in blocking solution (3% bovine serum albumin and 0.5% Triton X-100 in PBS) for 30 min at room temperature and then incubated in blocking solution containing goat anti-GFP (to recognize BFP, 1:1000, Abcam, ab6673) and rabbit anti-RFP (to recognize mCherry, 1:1000, MBL, PM005) antibodies over night at 4°C. Subsequently, sections were washed with PBS three times (10 min each) and incubated in blocking solution for 2 h at room temperature with fluorescent donkey anti-goat alexa fluor 488 (1:500, Invitrogen, A11055) and goat anti-rabbit alexa fluor 647 (1:500, Invitrogen, A21245). Finally, immuno-labelled sections were rinsed three times with PBS, mounted with an anti-fade mounting media (Solarbio, S2100) dehydrated and coverslipped.

For in situ hybridization in hypothalamic slices, brains were sectioned at 40 μm. (VT1000S, Leica). DNA templates for generating in situ probes were cloned using the Vgat primer: 5′-gccattcagggcatgttc-3′ and 5′-agcagcgtgaagaccacc–3′. Anti-sense RNA probes were transcribed with T7 RNA polymerase (Promega, Cat# P207E) and digoxigenin (DIG)-labeled nucleotides. As previously described^49^, sections were incubated with sheep anti-digoxygenin-POD (1:500, Roche, Cat# 11093274910) in 0.5% blocking reagent (Roche, 11096176001) for 24 h. After three washes in TBST, the reaction was in TSA-Cy5 (1:1000, Biotend, Cat# NEL745E001KT) diluted by TSA reaction buffer (1:100 volume mixture by 0.15% H_2_O_2_ and 20 times diluted 1M Tris-HCl pH=7.5) for 30 min before final washes in TBST for 3 times. For immuno-staining after in situ hybridization, the primary antibody rabbit anti-dsRed (Takara Bio, Cat# 632496, 1:500) and the secondary antibody goat anti-rabbit, Cy3 (Jackson ImmunoResearch, Cat# 111-655-003, 1:2,000) were used.

Images from cell cultures were captured using a Nikon Eclipse 80i fluorescence microscope. Images from brain slices were acquired by an Olympus FV3000 confocal microscope. For analysis of endogenous fluorescence intensities, hippocampal contours were drawn in Fiji. Mean fluorescence value of hippocampal slices were measured after background subtraction. Typically, 2 – 10 slices (FOV) per brain were analyzed. For co-labeling analysis, confocal images were imported into Imaris (v7.3, Bitplane) for cell counting.

### Western blot

The transfected HEK293T cells in 24-well-plate were lysated with 200μl Protien lysis buffer (50 mM Tris-HCl pH6.8, 2% SDS, 0.1% Bromophenol blue, 10% glycerol, 1% β-Mercaptoethanol), denatured for 10 min at 95 °C before loading. Using a 12 % sodium dodecyl sulfate (SDS) polyacrylamide gel to separate proteins, samples were transferred to a PVDF membrane (Millipore, #IPVH00010). The blotted membranes were blocked with blocking solution containing 5% non-fat milk powder (Sangon Biotech, A600669) in PBS-T (0.1% Tween-20 in PBS buffer) for 60 min, followed by an overnight exposure at 4 °C in the same buffer to the following primary antibodies: (a) rabbit anti-mCherry polyclonal antibody (1:5000, Millipore #AB356482); (b) rabbit anti-GFP polyclonal antibody (1:2000, Proteintech 50430-2-AP); (c) mouse antiβ-Tubulin monoclonal antibody (1:10000, ProteinTech # 66240-1-Ig). The membranes were washed and incubated for 2 h at room temperature with the corresponding secondary antibodies: (a) HRP-conjugated goat anti-rabbit IgG (1:10000, ZSGB-Bio #ZB-5301); (b) HRP-conjugated goat anti-mouse IgG (1:10000, ZSGB-Bio #ZB-5305). Signals were detected and visualized by exposure to Clarity Western ECL Substrate (BioRad, #1705061) and digitized with Tanon 5500 chemiluminescence imaging analysis system.

### Slice electrophysiology

Mice were first anaesthetized by isoflurane (5%) and then deeply anaesthetized by high-dose isoflurane (~150 μL in a custom nose mask). Mice were transcardially perfused with ice-cold NMDG-based slicing artificial cerebral spinal fluid (ACSF, 93 mM NMDG, 2.5 mM KCl, 1.2 mM NaH_2_PO_4_, 30 mM NaHCO_3_, 20 mM HEPES, 5 mM Sodium ascorbate, 2 mM thiourea, 3 mM Sodium pyruvate, 25 mM D-glucose,12 mM N-Acetyl-L-cysteine, 10 mM MgCl_2_, 0.5 mM CaCl_2_, oxygenated with 95% O_2_/5% CO_2_). After the perfusion, the brain was immediately removed and transferred to ice-cold NMDG-based slicing ACSF. Coronal brain slices (300 μm thick) containing hippocampus were prepared using a vibratome (VT-1200S, Leica) in an ice-cold NMDG-based ACSF. Slices were maintained for 12 min at 32°C in NMDG-based ACSF and were subsequently transferred into HEPES-based solution containing (in mM: 92 NaCl, 2.5 KCl,1.2 NaH_2_PO_4_, 30 NaHCO_3_, 20 HEPES, 5 Sodium ascorbate, 2 thiourea, 3 Sodium pyruvate, 25 D-glucose, 2 MgCl_2_, 2 CaCl_2_, bubbled with 95% O_2_/5% CO_2_) at room temperature and incubated more than 1 h before recording, and then were kept at room temperature (20-22°C) until start of recordings. For the recording, slices were transferred to a recording chamber and infused with ~30 °C recording ACSF (in mM: 119 NaCl, 5 KCl, 1.25 NaH_2_PO_4_, 26 NaHCO_3_, 10 D-glucose, 1 MgCl_2_, 2 CaCl_2_, bubbled with 95% O_2_ / 5% CO_2_). All chemicals were purchased from Sigma-Aldrich.

Patch pipettes (4 – 7 MΩ) pulled from borosilicate glass (Sutter instrument, BF150-86-10) were filled with a K-gluconate based internal solution (in mM: 126 K-gluconate, 2 KCl, 2 MgCl_2_,10 HEPES, 0.2 EGTA, 4 MgATP_2_, 0.4 Na_3_GTP, 10 Na-phosphate creatine, 290 mOsm, adjusted to pH 7.2~7.3 with KOH). Whole-cell recording was performed with a Multiclamp 700B amplifier and a Digidata 1440A (Molecular Device). Voltage-clamp recording was conducted at holding potential −70 mV. Current-clamp recording was conducted at membrane potential −70 mV. For optogenetic stimuli, the blue light pulse (480 nm) from a LED engine (Sola SE5-LCR-V8, Lumencor) was triggered by digital commands from the Digidata 1440A and delivered through the objective to illuminate the entire field of view. The yellow light from a DPSS laser (589 nm, Shanghai Century) was delivered through optical fiber positioned above the brain slice under the objective. The power at tip of fiber was ~18 mW with ~1 mm distance to the recording site.

To obtain the primate brain slices, the cynomolgus monkey was maintained under general anesthesia. The craniotomy was re-opened, and a fresh brain tissue block was cut from an injection site and immediately transferred to ice-cold NMDG-based ACSF. Primate brain slices were prepared similarly as mouse brain slices and recorded in recording ACSF. Animal was monitored continuously until the slice preparation and recording were done for all injected sites, and then was euthanized with an overdose tiletamine hydrochloride and zolazepam hydrochloride.

### Behaviors and photometric recording

Four to six weeks after stereotaxic injections, animals were implanted with 200 μm diameter optic fibers (NA 0.37, Anilab, China) in the ventral CA1 (dorsal part: AP −3.2mm, LAT 3.4mm, DV −1.2mm; ventral part: AP −3.2mm, LAT 3.4mm, DV −3.3mm). Animals were singly housed and were allowed one week for recovery after the surgery. Animals were habituated to the experimenter by handling for at least 5 days beforehand, and the behavioral experiments were conducted during the animal’s light cycle. For open field test, animals were placed in an open arena (40 × 40 × 45 cm) and allowed to explore for 15 min. For conditional place preference (CPP), the apparatus consists of two square contexts (25 × 25 × 30 cm, connected via a bridge 7 × 10 × 30 cm) with identical pump port on the opposite site but distinct walls and odors. Animals were water restricted and maintained at 90 % body weight two days before experiments. During habituation, mice were habituated to the two connected contexts for 15 min without water reward. During the training sessions (day 2 – 4), mice were first restricted in the rewarding context for 10 min (2 μl drop upon licking, one drop per 5 s), and then restrained in non-rewarding context for another 10 min (no water). On day 5, animals were allowed to freely explore the two contexts for 15 min without reward (CPP memory retrieval) and then for another 15 min with water reward available. On day 6, animals went through the same procedure as day 5 except that setup orientation was reversed and the reward was changed to sucrose. The CPP score is calculated as (A-B)/(A+B) where A and B are the time that the animal has stayed in the two contexts respectively. For fear conditioning, animals received six foot-shocks (2 s, 0.7 mA, 60 s interval) in the conditioning context.

The optic fibers were connected with a 465 nm blue LED light source (Cinelyzer, Plexon Inc). The excitation light at the tip of fiber was adjusted to be 14 – 16 μW (setting 10 – 12 in the LED controller). Emission signals and behavioral videos (640 × 480 pixels at 30 Hz) were captured simultaneously by a multi-fiber photometry recording system (Cinelyzer, Plexon Inc).

Raw data of emission signals were acquired and directly extracted for analysis in MATLAB (MathWorks) without any filtering. The area (mean) of the Ca^2+^ signals in a defined window were quantified for the event-related responses. The 0.5 s period before water/sucrose reward onset was computed as the baseline and the 3 s period after stimulus onset was computed as the response. The 2 s before foot shock onset was computed as the baseline and the 10 s period after stimulus onset was computed as the response. For polar plot, the averaged response for each type of projection cells was computed in each animal. Then the responses were averaged across animals in each projection type and normalized by the maximal response in all projection types. To analyze photometric signals when the animal is moving to the center of the open field (the locomotion is continuous), the filtered Ca^2+^ signal from the Cinelyzer was used (smoothed with an exponentially weighted moving average^50^, tau= 0.2 s). The center area is defined as a rectangle with 40% of width and length of the open arena. For each center entry trial, the 3 s before the enter entry was computed as the baseline and 3 s after the entry was computed as the signal changes related to the center entry. Short trials (<6 s) were not included for analysis.

### Quantification and statistical analysis

Mean values are accompanied by the standard error of the mean (S.E.M.). For the box plot, data are presented as median (center line) with 25/75 percentile (box) and min and max of the data points (whiskers); the plus or filled circle indicates mean. No statistical methods were used to predetermine sample sizes. Statistical analysis was performed in GraphPad Prism 6. Mann-Whitney U test, Wilcoxon matched-paired signed rank test, two-sided paired *t*-test, and one-way ANOVA test were used to test for statistical significance. Statistical parameters including the exact value of n, precision measures (mean ± SEM) and statistical significance are reported in the text and in the figure legends (see individual sections). The significance threshold was placed at *a* = 0.05 (n.s., P > 0.05; *, P < 0.05; **, P < 0.01; ***, P < 0.001; ****, P < 0.0001).

